# Transient RNA structures underlie highly pathogenic avian influenza virus genesis

**DOI:** 10.1101/2024.01.11.574333

**Authors:** Mathis Funk, Monique I. Spronken, Theo M. Bestebroer, Anja C.M. de Bruin, Alexander P. Gultyaev, Ron A.M. Fouchier, Aartjan J.W. te Velthuis, Mathilde Richard

## Abstract

Highly pathogenic avian influenza viruses (HPAIVs) cause severe disease and high fatality in poultry^1^. They emerge exclusively from H5 and H7 low pathogenic avian influenza viruses (LPAIVs)^2^. Although insertion of a furin-cleavable multibasic cleavage site (MBCS) in the hemagglutinin gene was identified decades ago as the genetic basis for LPAIV-to-HPAIV transition^3,4^, the exact mechanisms underlying said insertion have remained unknown. Here we used an innovative combination of bioinformatic models to predict RNA structures forming around the influenza virus RNA polymerase during replication, and circular sequencing^5^ to reliably detect nucleotide insertions. We show that transient H5 hemagglutinin RNA structures predicted to trap the polymerase on purine-rich sequences drive nucleotide insertions characteristic of MBCSs, providing the first strong empirical evidence of RNA structure involvement in MBCS acquisition. Insertion frequencies at the H5 cleavage site were strongly affected by substitutions in flanking genomic regions altering predicted transient RNA structures. Introduction of H5-like cleavage site sequences and structures into an H6 hemagglutinin resulted in MBCS-yielding insertions never observed before in H6 viruses. Our results demonstrate that nucleotide insertions that underlie H5 HPAIV emergence result from a previously unknown RNA-structure-driven diversity-generating mechanism, which could be shared with other RNA viruses.

---

Over the past 60 years, 48 independent HPAIV emergence events have been described^2^. Outbreaks of HPAIV have devastating impacts on domestic and wild bird populations and present a global health risk due to repeated spill-over events into mammals including humans^6^. HPAIV emergence always occurs following acquisition of an MBCS in the hemagglutinin (HA) gene, which encodes the influenza virus receptor-binding and fusion protein that mediates virus entry into cells. Translation of the HA gene yields a precursor protein which is post-translationally cleaved by host-cell proteases, a step required for activation of the fusion potential of HA and therefore virus infectivity^7^. While LPAIV HAs have a monobasic cleavage site that can only be cleaved by tissue-restricted trypsin-like proteases, HPAIV MBCSs are cleaved by ubiquitous furin-like proteases, allowing systemic virus spread in terrestrial poultry^2^. Although the key genetic basis of LPAIV-to-HPAIV conversion — i.e., MBCS acquisition — has been known since the 1980s^3,4^, the underlying mechanisms are still unknown. Genetic analysis of MBCSs from newly converted HPAIVs suggests three different origins. Two result from errors by the influenza virus RNA-dependent RNA polymerase (RdRp) during influenza virus RNA replication^2,8–11^. These errors are either nucleotide substitutions leading to non-basic to basic amino acid changes, and/or insertions of nucleotides coding for basic amino acids, probably by duplication of neighboring sequences. The third, observed in H7 only, results from non-homologous recombination with host or viral RNA^12–16^, in which the involvement of the RdRp is not yet clear. Here, we sought to elucidate the mechanism of MBCS acquisition by duplication, as it is the most likely mechanism underlying 36 of the 45 HPAIV conversion events with known MBCS sequences and the majority of H5 LPAIV-to-HPAIV conversions^2^.

It has been shown that the restriction of MBCS acquisition to H5 and H7 HAs is not due to incompatibilities at the protein level, since non-H5/H7 HAs can accommodate artificial MBCSs that can be cleaved by endogenous proteases^17–21^. Instead, we and others have hypothesized that insertions at the HA cleavage site in H5 and H7 HAs might be due to HA RNA sequence and/or structure^8,9,11,22–27^. The functional influenza virus replication unit is a ribonucleoprotein (RNP) complex comprising the RdRp (PB2, PB1 and PA subunits), nucleoprotein (NP) and a template RNA, containing highly conserved 5’ and 3’ promoter sequences. While vRNAs (negative-sense genomic influenza virus RNAs) and cRNAs (positive-sense copies of vRNA serving as replication intermediates) are bound by NP, the non-uniformity of NP coverage allows secondary RNA structures to form in cells and virions^28,29^. Perdue et al. hypothesized that the RdRp can be stalled during replication by downstream RNA structures, leading to duplications^9^. However, this model is incompatible with our previous observation that the insertions are located in the loop of conserved in silico predicted stem-loop structures^22^. Recent insights from RdRp structures during transcription revealed that the template entrance and exit channels are in close proximity^30,31^, theoretically allowing the formation of double-stranded RNA structures between the entering and exiting template strands. These structures would prevent movement of the template into the active site, impeding replication **(Fig. 1a)**, and would be transient since the entering and exiting RNA template brought into close proximity shift as the RdRp moves from the 3’ to the 5’ end of the template RNA. By combining this new knowledge and the prediction of stem-loop RNA structures at the HA cleavage site, we have proposed a novel model: the trapped RdRp model^2,25^. In this model, a transient RNA structure is formed by the template during RNA replication, enclosing and trapping the RdRp. This physical constraint on the RdRp causes it to stutter (replicate the same nucleotide repeatedly) or backtrack (sliding back along the template over several nucleotides before reinitiating replication), causing an increase in duplication insertions. Previous studies have highlighted that viruses with adenine/uracil-(A/U)-rich HA cleavage sites are more prone to accumulate insertions and convert to HPAIV^23,27,32–38^, indicating that sequence of the HA cleavage site also plays an important role. As conserved stem-loop RNA structures were also predicted at the HA cleavage site of non H5/H7 viruses^22,25^, their presence alone may not be sufficient for insertions. Therefore, we further hypothesize that the RdRp needs to be trapped by transient RNA structures on A/U-rich sequences to generate duplications, and that both factors act synergistically. In another context, putative RdRp-trapping structures have recently been shown to interfere with the replication of short aberrant influenza virus RNAs^39^, supporting the previously proposed model for insertions at the HA cleavage site^2,25^.

**Fig. 1:**
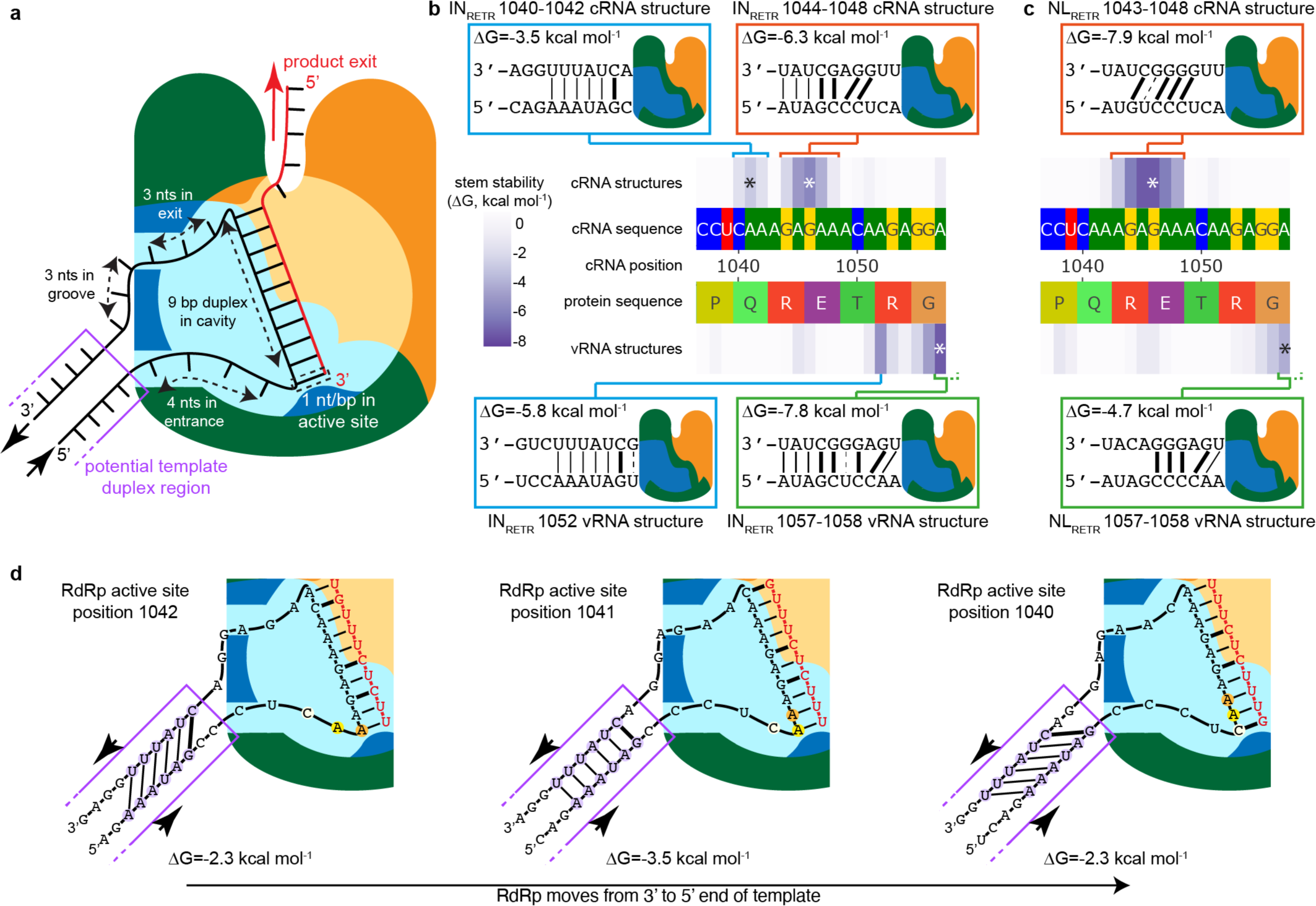
The trapped RdRp model and predicted transient RNA structures at the HA cleavage site. **a**, Schematic representation of the RdRp (PB2 in orange, PB1 in blue, PA in green), during replication of a template RNA (black) into a product RNA (red). The template threads through the RdRp (approximate lengths based on PDB6SZU^44^) and exits adjacent to the template entrance, allowing potential base pairing. Nt: nucleotide, bp: base pair. **b-c**, Predicted RdRp-trapping structures at the HA cleavage sites of A/Indonesia/5/2005 (IN_RETR_, **b**) and A/mallard/Netherlands/3/1999 (NL_RETR_, **c**). The nucleotide sequence of the HA cleavage site (cRNA sense) and the corresponding amino acid sequence are colored by nucleotide or amino acid. Numbering starts at the first 5’ nucleotide of the segment. For each RdRp active site position, the free energy change ΔG of the RdRp-trapping structure is represented by the heatmaps according to the scale on the left. Lower ΔG values reflect higher RNA structure stability. Each transient RNA structure was named after the range of RdRp active site positions for which identical basepairs were predicted to form. Structures only identified in IN_RETR_ are shown in blue frames while structures predicted to form for analogous RdRp active site positions in both IN_RETR_ and NL_RETR_ are shown in orange (cRNA) and green (vRNA) frames. Brackets on the heatmap indicate that the same structure is predicted to form for several RdRp active site positions. The asterisk identifies the RdRp active site position when the structure is at its most stable conformation, corresponding to the schematic representation. A thick line indicates a G-C, a thin line an A-U, and a dashed line a G-U base pair. **d,** Detailed representation of the same cRNA structure predicted to form for RdRp active site positions 1040-1042 on IN_RETR_ cRNA, using the same representation as **a**.

The difficulty to reproduce the rare event of MBCS acquisition in laboratory settings has hampered the advancement of knowledge. In addition, the presence of highly repetitive sequences in MBCSs complicates the reliable detection of insertions, as these sequences are prone to insertions or deletions for many replication enzymes, such as reverse-transcriptases and DNA polymerases needed in the process of RNA sequencing. Here, we used a novel approach in which a virus-free replication system was combined with a customized circular sequencing^5^ approach to accurately identify and quantify insertions introduced by the RdRp during RNA replication. A virus-free system allows the characterization of the full RdRp insertional landscape free from protein translation and viral selection constraints. Our results show that two factors directly influence insertion nature and frequency at the H5 cleavage site: the formation of putative RdRp-trapping RNA structures and the template RNA sequence in the RdRp active site cavity, with sequences containing long A stretches being more insertion-prone.

## Predicting RdRp-trapping RNA structures

Since it is well-established that insertions in H7 HAs often occur via non-homologuous recombination rather than potential duplication^2^, we focused on investigating the molecular mechanism underlying insertions at H5 HA cleavage sites. We chose two model sequences: A/mallard/Netherlands/3/1999 HA (NL_X,_ where X is the amino acid sequence of the HA cleavage site) and A/Indonesia/5/2005 HA (IN_X_). The former was chosen as its cleavage site nucleotide sequence, as well as the 30 nucleotides up- and downstream, correspond to the consensus of viruses from the Africa-Eurasia-Oceania (AEO) H5 LPAIV lineage^40^. The latter is a representative of H5 HPAIVs of the A/goose/Guangdong/1/1996 (Gs/Gd) lineage, which first emerged in 1996 and is still circulating in birds^41–43^. HAs of Gs/Gd lineage viruses have acquired substitutions in regions flanking the HA cleavage site that stabilize predicted RNA structures^22^. To mimic an LPAIV HA, the wild type (WT) IN_RESRRKKR_ MBCS was removed and replaced with the highly conserved H5 LPAIV cleavage site nucleotide sequence, coding for the amino acid sequence RETR.

To investigate the impact of putative transient RNA structures on insertions, we designed an RdRp-trapping structure prediction tool. We modelled transient RNA structures instead of static structures as might exist in non-replicating vRNPs^22,25,27^ by using a sliding window approach as described previously^39^. The 20-nucleotide RdRp footprint^44^, corresponding to the RNA that is interacting with the RdRp, was slid along the RNA template sequence nucleotide per nucleotide, in both cRNA and vRNA sense. For each nucleotide in the RdRp active site, referred to as RdRp active site positions hereafter, the free energy change (ΔG) of RNA structure formation between 10-nucleotide-long RdRp-flanking regions on either side was determined while mimicking the close proximity of entering and exiting template strands (see **Methods**, **Extended Data Fig. 1)**. Predicted RdRp-trapping RNA structures at the HA cleavage site of IN_RETR_ and NL_RETR_ are shown in **Figure 1b,c**, and an interactive graph showing predictions for all HA sequences used in this manuscript is provided as **Supplementary Fig. 1**. When predicted transient RNA structures contain unpaired nucleotides close to the RdRp, identical base pairs can form for different RdRp active site positions (**Fig. 1d**). Differences in the number of unpaired nucleotides on each side of the RdRp might still lead to varying predicted stability, with symmetrical distributions being more stable.

Four main transient RNA structures were predicted to form in IN_RETR,_ while only two of them were predicted in NL_RETR_. The structures found only in IN_RETR_ result from the aforementioned Gs/Gd lineage-specific substitutions^22^, absent in H5 LPAIV HAs like NL_RETR_. The shared cRNA structure predicted to form for RdRp active site positions 1043/1044-1048 for both IN_RETR_ and NL_RETR_ was more stable in NL_RETR_, while the shared predicted vRNA structure for RdRp active site positions 1057-1058 was more stable in IN_RETR_. Importantly, MBCS-yielding insertions have occured in nature between positions 1041-1052^2^, encompassing the IN-specific structures and the shared cRNA structure.

## Insertions are promoted by A/U stretches

To experimentally assess insertion frequencies observed at the HA cleavage site, we generated HA RNPs via transfections in two biological replicates, and vRNA-sense HA extracted from cells was sequenced using a custom circular sequencing method^5^. Here we focused on insertions, as they are the most relevant for H5 MBCS acquisition. Insertion frequencies were corrected for non-RdRp background insertions and potential duplications were realigned to the most stable predicted RdRp-trapping structure, standardizing insertion placement and testing compatibility with our model (see **Supplementary Note 1**, **Methods**). We first investigated the impact of HA cleavage site sequence on insertions. Virtually no insertions were observed at the LPAIV IN_RETR_ and NL_RETR_ HA cleavage site, though a few instances of one- or two-A insertions were observed (**Fig. 2a**). We and others have previously suggested that H5 MBCS acquisition occurs following initial nucleotide substitutions in the genetically stable LPAIV RETR cleavage site^23,36,40^. The RETR-encoding cleavage site sequence was threrefore replaced by a sequence encoding REKR, suggested previously to be a stepping-stone for MBCS acquisition^34,36^, or RKKR, shown previously to be insertion-prone^23,27,32,45^ (all HA nucleotide sequences are provided as **Supplementary Data 1**). The REKR cleavage site sequence led to increased, but still very low insertion frequencies (IN_REKR_: 0.8-1.1‰, NL_REKR_: 0.8-0.9‰, **Extended Data Fig. 2a**), while insertions were detected at much higher frequencies in RKKR-bearing HAs (**Fig. 2b**), up to 30.5‰ for GA insertions at position 1044 (1044GA) in NL_RKKR_. The insertions observed in the REKR- and RKKR-containing HAs were highly reminiscent of those seen in H5 MBCSs in nature, consisting of additional As and Gs between cRNA positions 1041-1052^2^ (**Supplementary Fig. 2**). Insertions that could be explained by a single duplication were sorted into three types: 1) single-nucleotide insertions, most commonly 1046A, 2) homopolymer insertions, most commonly 1046AA or 1046AAA, and 3) heteropolymer insertions, most commonly 1044GA or 1044GAA. The most common insertions for all three groups occurred in or adjacent to the 1045-1052 eight-A stretch. A minority of insertions could not be explained by direct duplications of adjacent sequences and were sorted into a fourth type: complex insertions (IN_RKKR_: 0.6-0.6‰, NL_RKKR_: 1.7-2.0‰). They may be explained by several consecutive or concomitant insertion, deletion, and/or substitution events. Some complex insertions also most likely resulted from non-homologuous recombination (**Supplementary Note 2)**. Uncorrected insertion frequencies for each category for each template and replicate can be found in **Supplementary Data 2.**

**Fig. 2:**
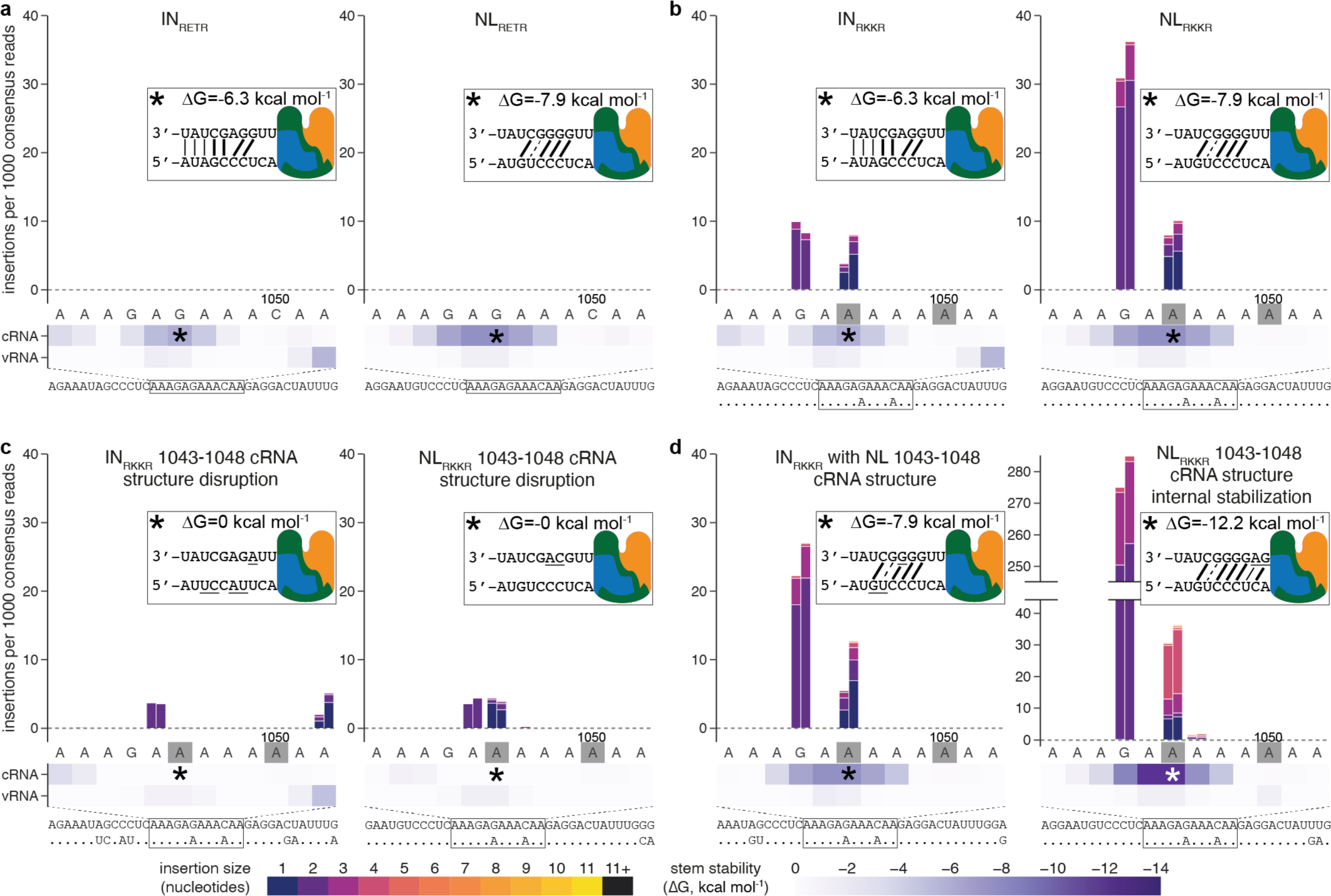
The predicted 1043-1048 cRNA structure results in insertions in A/U-rich cleavage sites. **a-d,** Insertion frequencies observed in A/Indonesia/5/2005 (IN, left) and A/mallard/Netherlands/3/1999 (NL, right), with the consensus H5 LPAIV RETR cleavage site (**a**), with an RKKR cleavage site (**b**), with an RKKR cleavage site and substitutions disrupting the 1043-1048 cRNA structure (**c**), with RKKR cleavage site and substitutions stabilizing the 1043-1048 cRNA structure (**d**). Stacked bars show the number of insertions per 1000 consensus reads, background-corrected by subtracting frequencies observed in non-RdRp samples. The color of the bar reflects the size of the insertion, according to the bottom legend. Results of two independent replicates are shown. The cRNA sense nucleotide sequence is indicated on the x-axis. Heatmaps below the x-axis indicate the stability (ΔG, expressed as kcal mol^-1^) of predicted RdRp-trapping structures for each RdRp active site position, according to the bottom heatmap scale. The sequence below the graphs shows all substitutions relative to the RETR-containing HA. Insets in each graph show a schematic representation of the RdRp-trapping stucture predicted to form for the RdRp active site position marked by an asterisk on the heatmap, using the same representation as in **Fig. 1**, with substitutions relative to the corresponding RETR-containing HA underlined. Non-corrected insertion frequencies for RdRp-containing and background samples are provided as an interactive figure in Supplementary Fig. 2.

Heteropolymer insertions were more common than single-nucleotide insertions and homopolymer insertions in NL_RKKR_ (31.0-36.3‰, 4.6-5.3‰, 2.2-3.3‰, respectively) and IN_RKKR_ (8.5-10.1‰, 2.4-5.1‰, 1.0-2.5‰, respectively). Heteropolymer frequencies were found to vary little across 21 independent IN_RKKR_ biological replicates or when using different primers for RNA sequencing, while frequencies of other insertion types were more variable (**Supplementary Note 3**, **Extended Data Fig. 3a,b**). Insertions of two nucleotides were the most frequent, and frequencies declined steeply as sizes increased beyond two nucleotides (**Extended Data Fig. 3d**). As a result, only 1.6-2.1‰ and 5.1-7.3‰ consensus reads presented in-frame insertions for IN_RKKR_ and NL_RKKR_ respectively (**Supplementary Fig. 3**). The most common in-frame insertions for both HAs were 1044GAA heteropolymer insertions, resulting in an MBCS found in several newly converted HPAIVs in nature^2^ (AGAAGAAAAAAAGA; RRKKR). Across all samples, background insertions were almost exclusively insertions of 1 A, consistent with DNA polymerase I (PolI) erorrs^46^ (**Supplementary Note 4**).

## Insertions are driven by the 1043-1048 cRNA structure

Most insertions detected in IN_RKKR_ and NL_RKKR_ were consistent with potential duplications following RdRp-trapping by the transient cRNA structure predicted for RdRp active site positions 1043/1044-1048, referred to as the 1043-1048 cRNA structure hereafter. In addition, NL_RKKR_ led to higher insertion frequencies than IN_RKKR,_ suggesting that this shared cRNA structure, more stable in NL, might play a more important role than the IN-specific predicted RNA structures.

To assess the role of the 1043-1048 cRNA structure in driving insertions at the HA cleavage site, mutated HAs were designed in which this structure was specifically disrupted via 7 and 2 nts substitutions in the regions flanking the IN_RKKR_ and NL_RKKR_ cleavage sites, respectively. In parallel, we designed mutated HAs with 13 or 10 substitutions, respectively, disrupting all predicted transient RNA structures. Both the targeted (**Fig. 2c**) and the full disruption (**Extended Data Fig. 2b**) approach led to a decrease in heteropolymer insertion frequencies of, respectively, two- to threefold in IN_RKKR_ (8.5-10.1‰ to 3.7-3.9‰ and 3.0-3.4‰), and over eight- and fivefold in NL_RKKR_ (31.0-36.3‰ to 3.7-4.6‰ and 5.5-6.8‰). This suggested that the 1043-1048 cRNA structure plays an important role for heteropolymer insertions, but that there is a basal heteropolymer insertion frequency in RKKR-bearing cleavage sites in absence of predicted transient RNA structures. Complex insertion frequencies decreased about fourfold for IN_RKKR_ (0.6-0.7‰ to 0.1-0.3‰) and about sixfold for NL_RKKR_ (1.7-2.2‰ to 0.2-0.3‰), but single-nucleotide and homopolymer insertion frequencies showed a less consistent trend, either being stable or decreasing when predicted RNA structures were disrupted (**Supplementary Note 5**). Sorting insertions by size instead of type revealed that larger insertions — composed of heteropolymer, homopolymer and complex insertions — were more affected by disruption approaches.

Next, we stabilized predicted transient RNA structures. We introduced 3 substitutions flanking the IN_RKKR_ cleavage site predicted to change the IN 1043-1048 cRNA structure to the more stable NL one. Heteropolymer insertions frequencies increased by two- to threefold (8.5-10.1‰ to 22.3-27.1‰), to levels closer to NL_RKKR_ (31.0-36.3‰, **Fig. 2d**). Single-nucleotide insertion frequencies showed little variation (2.4-5.1‰ to 2.5-6.7‰), but homopolymer insertions frequencies showed a twofold increase (1.0-2.5‰ to 2.3-4.7‰). Splitting insertions by size showed that two-nucleotide insertions increased about twofold (9.1-9.6‰ to 19.8-25.0‰), while larger insertions increased about threefold (1.9-2.4‰ to 5.7-8.2‰). Stabilization of only the IN-specific 1040-1042 cRNA/1052 vRNA structures did not lead to an increase in insertion frequencies (**Extended Data Fig. 2e**).

To further corroborate the role of the 1043-1048 cRNA structure in driving heteropolymer and large insertions, we modified NL_RKKR_ in two ways predicted to stabilize this structure. First, the length of the double-stranded region was increased by two substitutions creating two additional base pairs at the edge most distant from the RdRp. This did not lead to an appreciable change in insertion frequencies (e.g., heteropolymer: 31.0-36.3‰ to 27.6-29.6‰, **Extended Data Fig. 2f**). The predicted 1043-1048 cRNA structure was then elongated with two additional base pairs at the edge closest to the RdRp, hypothetically tightening it around the RdRp. This resulted in an eightfold increase in heteropolymer insertion frequencies (31.0-36.3‰ to 275.2-285.3‰), with about 250.3-257.3‰ of consensus reads presenting the 1044GA heteropolymer insertion (**Fig. 2d**). There were also many more complex insertions (1.7-2.0‰ to 32.2-36.6‰), with notably 16.0-19.1‰ of consensus reads presenting a 1046GAGA insertion (0.2-0.2‰ in WT NL_RKKR_), likely the result of two 1044GA heteropolymer insertions. Single-nucleotide insertion frequencies decreased five- to tenfold (4.6-5.3‰ to 0.9-1.2‰), while homopolymer insertion frequencies remained within a twofold range (2.2-3.3‰ to 1.6-1.9‰). Both two-nucleotides and larger insertions increased about eightfold (28.5-33.1‰ to 252.6-259.4‰ and 6.0-8.1‰ to 49.0-57.2‰, respectively). Taken together, these results showed that, while single-nucleotide insertions might be primarily affected by the sequence in the RdRp active site, heteropolymer and large insertions observed in RKKR-bearing H5 HAs are affected by changes in the predicted 1043-1048 cRNA RdRp-trapping structure.

## Heteropolymer insertions occur on cRNA

Since the 1043-1048 cRNA structure was predicted to form only in cRNA sense, we hypothesized that heteropolymer insertions and insertions of more than two nucleotides occur when a cRNA template is replicated into a vRNA product by the RdRp. We therefore developed a unidirectional RNP replication system (see **Methods**, **Extended Data Fig. 4**). Insertion frequencies were not background-corrected since different RNA senses were targeted in RdRp-containing and non-RdRp samples leading to differences in observed insertion frequencies (see **Methods, Supplementary Note 6**). When only cRNA to vRNA replication occurred, similar insertion frequencies were observed compared to a bidirectional system (e.g., heteropolymer: 18.4-20.5‰ to 15.9-18.0‰, **Fig. 3a,b**). In contrast, when cRNA to vRNA replication was blocked, heteropolymer insertion frequency was strongly decreased compared to the bidirectional system (10.6-10.9‰ to 0.7-2.2‰, **Fig. 3c,d**). Additionally, frequency of complex insertion decreased over twofold when cRNA to vRNA replication was blocked (1.8-1.9‰ to 0.2-0.7‰), but not those of single-nucleotide (28.9-32.2‰ to 16.1-23.0‰) or homopolymer (4.7-7.7‰ to 3.1-5.4‰) insertions. Both two-nucleotide and larger insertions decreased threefold (13.0-14.9‰ to 4.3-4.5‰ and 3.5-5.1‰ to 1.1-2.0‰). Taken together, these results showed that heteropolymer insertions at the IN_RKKR_ cleavage site mainly occur during cRNA to vRNA replication and further confirm the role of the 1043-1048 cRNA structure in driving heteropolymer and large insertions at the HA cleavage site. Of note, the 1043-1048 cRNA structure was also highly conserved in HAs of putative LPAIV precursors of HPAIV outbreaks **(Supplementary Note 7**).

**Fig. 3:**
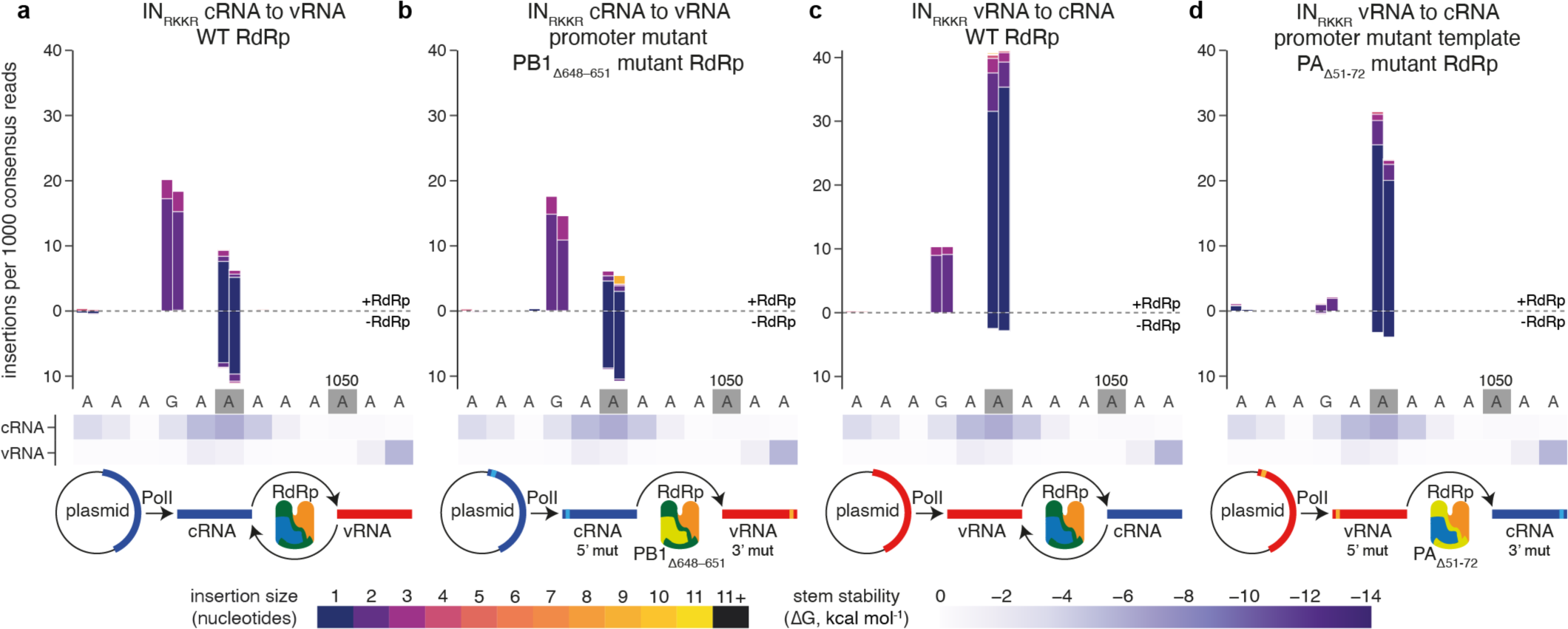
Heteropolymer insertions occur during replication of cRNA templates. **a-d** Insertion frequencies observed in IN_RKKR_ vRNA in a bidirectional (**a**) or unidirectional (**b**) replication system, or in IN_RKKR_ cRNA in a bidirectional (**c**) or unidirectional (**d**) replication system. Using the same representation as Fig. 2. Uncorrected frequencies are shown, growing upward for RdRp-containing samples and downward for input (non-RdRp) samples. In input samples, cRNA (**a, b**) or vRNA (**c, d**) is targeted for sequencing. Schemes below the graphs show the directionality of replication in each assay.

## H6 cRNA structure lead to low heteropolymer insertion frequencies

HPAIV emergence has been restricted to viruses of the H5 and H7 subtypes^2^. The reasons for this subtype-restriction are still unknown, though we and others have suggested that it might be due to codon use at the HA cleavage site favoring substitutions to insertion-prone sequences in H5 and H7 HAs^18,21,23,40^. To assess the potential impact of transient RdRp-trapping structuress in non-H5/H7 HAs, we chose to study H6. H6 HAs are phylogenetically most closely related to H5 HAs and H6 LPAIV also circulate extensively in poultry^47^. The A/mallard/Sweden/81/2002 HA (SW) was used, as its IETR cleavage site corresponds to the H6 LPAIV nucleotide consensus. Additionally, SW presents a predicted RdRp-trapping cRNA structure spanning RdRp active site positions 1038-1043, equivalent to positions 1043-1048 in H5 HAs (H5 numbering will be used hereafter). This structure is almost identical to the NL 1043-1048 cRNA structure, but with an A-T instead of G-C base pair at the third position. RNPs consisting of the H5 A/Indonesia/5/2005 RdRp and NP, and H6 SW_IETR_ RNA were produced, and only single-nucleotide insertions at extremely low frequencies were detected (0.0-0.0‰, **Fig. 4a**), similar to IN_RETR_ (0.0-0.0‰) and NL_RETR_ (−0.1-0.0‰). After replacing the IETR cleavage site with the RKKR sequence, heteropolymer insertion frequency increased to 4.2-5.0‰ (**Fig. 4b**), levels lower than in IN_RKKR_ and NL_RKKR_, but comparable to the basal heteropolymer insertion frequencies in RKKR-containing H5 HAs with disrupted structures (ranging between 1.8-6.8‰). Accordingly, disrupting the predicted H6 1043-1048 cRNA structure by introducing two substitutions did not reduce insertion frequencies, with all insertion types varying less than twofold (e.g., heteropolymer frequencies 4.2-5.0‰ to 4.0-5.4‰, **Fig. 4c**). However, stabilization of the predicted H6 1043-1048 cRNA structure, by introducing the NL G-C base pair in the third position of the structure, increased heteropolymer insertions about fivefold (4.2-5.0‰ to 20.7-20.9‰, **Fig. 4d**), levels closer to those observed in NL_RKKR_ (31.0-36.3‰). As observed previously, single-nucleotide frequencies showed little variation (4.2-4.7‰ to 4.5-8.2‰), while homopolymer insertion frequencies increased twofold (1.4-1.5‰ to 2.6-3.3‰). The increase was more pronounced for large insertions, with two-nucleotide insertions increasing fourfold (5.3-6.2‰ to 21.4-21.9‰) and larger insertions increasing sixfold (0.5-0.5‰ to 2.5-3.1‰). These results suggest that, in addition to the codon use at the cleavage site, predicted RdRp-trapping structures play an important role in the potential for HPAIV-yielding insertions in non-H5 HAs. Nevertheless, analysis of a set of 2070 unique H6 sequences^40^ showed that about 3% of H6 HAs possess the full NL 1043-1048 cRNA structure, suggesting that the main aspect preventing H6 HPAIV emergence may be the codon use at the cleavage site, disfavoring acquisition of an insertion-prone sequence^40^.

**Fig. 4:**
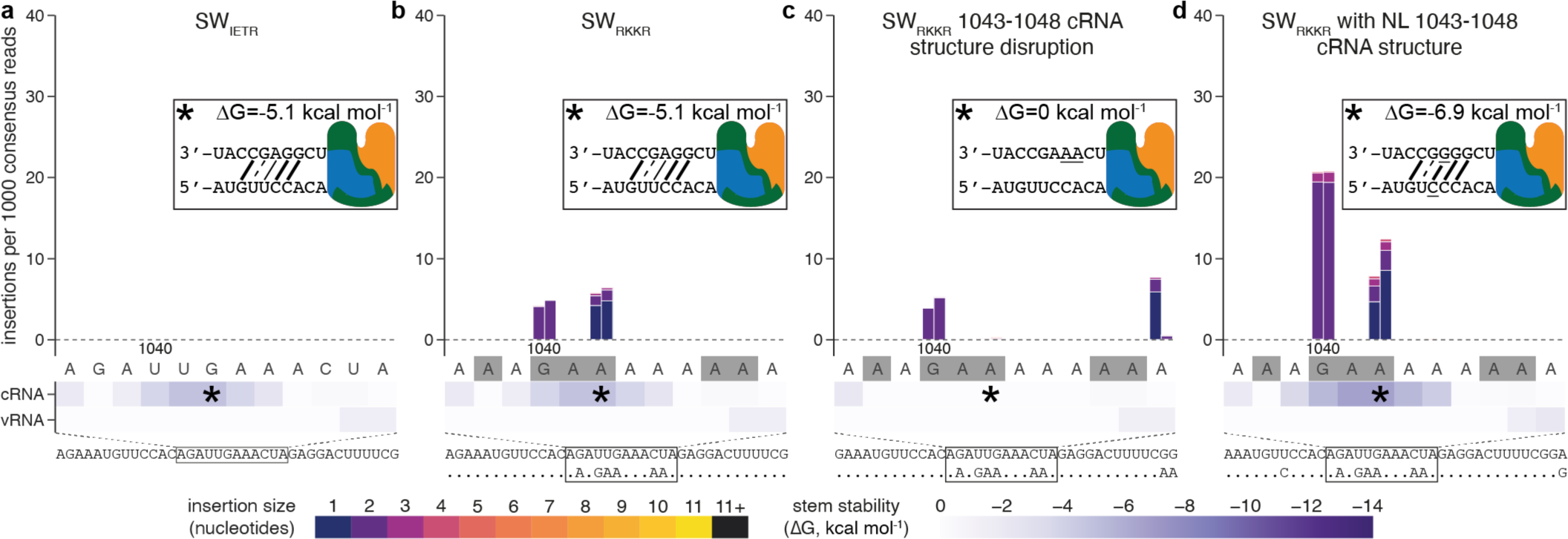
The cRNA structure of H6 A/mallard/Sweden/81/2002 HA (SW) induces less heteropolymer insertions than the H5 NL 1043-1048 cRNA structure. **a-d**, Insertion frequencies observed in the consensus H6 IETR cleavage site (**a**), with an RKKR cleavage site (**b**), with an RKKR cleavage site and substitutions disrupting the predicted 1043-1048 (H5 equivalent position) cRNA structure (**c**), with an RKKR cleavage site and substitutions stabilizing the predicted 1043-1048 cRNA structure (**d**). Using the same representation as **Fig. 2**.

## Discussion

The mechanisms underlying recurrent insertions at the H5 HA cleavage site have remained unsolved for decades. Whereas the importance of the HA cleavage site sequence has become increasingly supported^23,27,32–38^, we provide here for the first time empirical evidence of the involvement of RNA structures. While our data do not provide direct proof for transient RdRp trapping structure formation, model-guided stabilization or disruption of the 1043-1048 cRNA structure by only two nucleotide substitutions led to eightfold increase or decrease of heteropolymer insertions respectively, strongly supporting our model. A detailed animation showing how the most commonly observed insertion, 1044GA, could be explained by the trapped-RdRp model is provided as **Supplementary Video**.

Most insertions observed at the HA cleavage site could be the result of a single duplication event. Other insertions most likely reflected several successive duplication and/or deletion events or, very rarely, non-homologuous recombination, possibly via strand breakage in vRNPs and rejoining by cellular enzymes, as has also been suggested for pestiviruses^48–50^ and poliovirus^51^. Different types of putative duplication insertions showed different sensitivities to transient RNA-structure-affecting substitutions. Heteropolymer insertions, likely resulting from backtracking events, were strongly affected by transient RNA structures. Our data therefore suggest that backtracking, consisting of the RdRp sliding back several nucleotides on the template and duplicating several nucleotides in one event, requires physical trapping of the RdRp. On the other hand, single-nucleotide insertions, which likely occur following stuttering, i.e., repeated replication of the same template nucleotide, showed low sensitivity to transient structure-affecting substitutions. Notably, homopolymer insertions, which could result either from one or several backtracking or several stuttering events, exhibited variability in sensitivity to transient structures. While disruption of the 1043-1048 cRNA structure decreased homopolymer insertion frequencies in NL_RKKR_, it had little effect in IN_RKKR_. This discrepancy might be due to different relative contributions of structure-dependent backtracking and structure-independent stuttering events. Complex insertions most likely result from a heteropolymer insertion combined with another insertion or deletion and accordingly showed high sensitivity to both 1043-1048 cRNA structure stabilization and disruption. Overall, large heteropolymer, homopolymer and complex insertions were more senstitive to susbtitutions affecting the 1043-1048 cRNA structure, suggesting that they were more dependent on RdRp-trapping. Using unidirectional replication systems, we also showed that heteropolymer insertions were mainly generated during cRNA to vRNA replication, corroborating the important role of the 1043-1048 cRNA structure. This also confirms that heteropolymer insertions at the H5 cleavage site indeed occur on templates containing purine rather than pyrimidine stretches, as has been commonly assumed^9,23,27^. Deletions at the HA cleavage site showed sensitivity to A/U-richness, but contrarily to insertions showed very little sensitivity to transient RNA structure changes (**Supplementary Notes 8-10**).

Here we observed that insertions at the H5 HA cleavage site are exceedingly rare unless subsitutions are introduced changing the LPAIV RETR cleavage to a more insertion-prone sequence. The frequency of in-frame insertions that might lead to functional HA proteins remained very low, in line with the rarity of HPAIV conversions in nature. In-frame insertions observed here led to cleavage sites that were strikingly similar or even identical to H5 HPAIV MBCS sequences observed in nature (RRKKR, RKRKKR, RRRKKR). In fact, starting from a substituted cleavage site sequence (REKR, RKTR or RKKR), ten of the eleven MBCSs with insertion of known H5 HPAIVs can be explained by insertions observed in our system. They require either a single heteropolymer occurrence for six of them, or several consecutive heteropolymer or combinations of hetero- and homopolymer insertions for another four. If MBCSs can result from several consecutive insertion events occurring on the same molecule, frameshifting insertions could also play a role in MBCS acquisition. While the need for several insertions on the same molecule to restore the frame reduces the odds of occurence, additional insertions may have higher chances of accumulating in templates already bearing insertions leading to a more repetitive nucleotide cleavage site sequence. Only the MBCS of the H5 HPAIV responsible for the 1961 outbreak in South Africa outbreak seems unlikely to have occurred solely by duplication.

The exact requirements for transient structures to impede influenza RdRp replication have to be refined. The effect of transient structures may depend on both quantitative parameters, such as structure stability or RNA folding kinetics, and more qualitative parameters, such as structure tightness around the RdRp, as well as perhaps the nature of the base-pair forming closest to the RdRp. The position of the RNA structure in relation to the sequence present in the RdRp active site also plays a crucial role, as illustrated by the fact that stabilizing the 1043-1048 cRNA structure increased insertions, while stabilizing the IN-specific 1040-1042 cRNA structure did not. Additionally, the NP-binding profile of viral RNAs might affect RdRp-trapping structure formation. Recent work has shown that NP-binding profiles differ depending on influenza virus genomic sequence^28,52^. NP-bound RNA is thought not to be able to form secondary structures^53^. It is however important to note that during replication, NP molecules are dislodged from the template^44^, theoretically allowing formation of transient RNA structures between the (transiently) naked RNA strands protruding from the template entry and exit channels. Properties of the RdRp itself might also influence the sensitivity to RdRp-trapping structures^39,54,55^.

Along with providing a mechanism of insertions at the H5 HA cleavage site, our study provided new insights regarding the subtype specificity of HPAIV emergence. Introducing the insertion-prone RKKR sequence into an H6 HA, requiring 6 substitutions and highly unlikely to occur in nature^40^, only led to low frequencies of insertions, probably reflecting basal insertion levels in an A-rich sequence. Replacing the weaker H6 cRNA structure with the H5 cRNA structure increased insertions to ranges more comparable to those in insertion-prone H5 cleavage sites. This highlights the synergistic need for insertion-prone sequences and insertion-enhancing transient RNA structures and suggests that subtype specificity of HPAIV emergence might be explained by the combined presence of these factors, with perhaps an additional role played by the RdRp. While we focused here on insertions in H5 cleavage sites, most H7 HPAIV MBCS acquisitions also occurred following duplication-like insertions. While RNA sequences and structures found at H7 LPAIV cleavage sites differ markedly from those found in H5 LPAIVs^22,40^, a similar structure-driven mechanism might be involved.

The mechanism of insertions at the HA cleavage site unravelled here is reminiscent to those proposed in very different influenza virus contexts. Physical constraints on the template as it moves through the RdRp have been proposed to induce stuttering by repeated template-product realignment in the context of influenza virus mRNA polyadenylation^44^. Here too, an insertion-prone A or U stretch in the template RNA is required^56^. Putative transient RdRp-trapping RNA structures have recently also been shown to interfere with the replication by the RdRp of short aberrant influenza virus RNAs^39^. Replication of these short templates led to RdRp stalling in vitro or the formation of large internal deletions in cell culture. The replication errors occurred at positions consistent with predicted RdRp-trapping RNA structures and were sensitive to structure-affecting substitutions. In vitro replication experiments also showed that template RNAs remain bound by trapped RdRps, while product RNAs can dissociate from the RdRp-template complex, as would be required for backtracking insertions. RdRp replication errors induced by RNA structures might therefore be a more widespread phenomenon with different effects depending on yet to be elucidated factors, such as perhaps length of the template, presence of NP, or current replication state of the RdRp. This mechanism may thus also be involved in the recurrent insertions and deletions observed in the 160 antigenic loop of human influenza B viruses^57,58^ for example. Beyond influenza viruses, transient RdRp-trapping structures may also impact replication of other negative-sense RNA viruses with similar polymerase entry and exit channel architecture, such as Ebola virus, Lassa virus, measles virus or rabies virus^59^.

## Methods

### Plasmids

The generation of pCAGGS expression plasmids for A/Indonesia/5/05 PB2, PB1, PA and NP was described previously^60^. Plasmids expressing the mutated PB1 and PA genes were obtained by primer-directed PCR mutagenesis as described previously^61^. The generation of modified pHW2000 plasmids expressing genomic vRNAs of the HA segment of A/mallard/Netherlands/3/1999 (NL), A/mallard/Sweden/81/2002 (SW), and A/Indonesia/5/2005 (IN), whose MBCS was replaced by the LPAIV RETR cleavage site, under the control of the human DNA polymerase I promoter were described previously^62–64^. Mutated HA templates were subsequently generated by primer-directed PCR mutagenesis^61^. For the unidirectional RdRp replication systems, the IN_RKKR_ segment was transferred, in both vRNA and cRNA sense, to a pSP72 vector under control of the human DNA polymerase I promoter^62^ by “megaprimer” PCR mutagenesis, as described previously^65^. Influenza promoter substitutions were then introduced into these plasmids via primer-directed PCR mutagenesis^61^. Primers were ordered from Eurogentec. All plasmids were sequenced using a 3130XL or 3500XL Genetic Analyzer (Applied Biosystems) to confirm introduced mutations.

### Cells and generation of influenza virus RNPs

Human embryonic kidney cells (293T, ATCC) were cultured and transfected using CaCl_2_ as described previously^62^. To generate RNPs, 293T cells were transfected using 2.5 µg of plasmid encoding each A/Indonesia/5/2005 RdRp subunit (PB2, PB1, PA), 10 µg of plasmid encoding A/Indonesia/5/2005 NP, 5 µg of plasmid expressing HA cRNA or vRNA and 7.5 µg of empty pCAGGS plasmid. For background samples, the RdRp-encoding plasmids were omitted and an additional 7.5 µg of empty pCAGGS plasmid added.

### RNA extraction, fragmentation, and circularization

RNAs were extracted 48 hours after RNP transfection using the High Pure RNA Isolation Kit (Roche) according to instructions of the manufacturer. DNA was removed by performing 1 hour digestion with TURBO DNAse with subsequent bead clean up (Invitrogen) according to manufacturer’s instructions. 15-30 µL of RNA, depending on RNA concentration, were concentrated to 7.5 µL using the Oligo Clean & Concentrator Kit (Zymo Research) and fragmented by alkaline hydrolysis by adding 1 volume of 100 mM bicarbonate (Sigma-Aldrich) pH 9.2 buffer and incubating at 95 ℃ for 1-20 minutes depending on initial RNA concentration. After adding 1 volume of Gel Loading Buffer II (Invitrogen), RNA was run on a pre-warmed denaturing 8 M urea (Sigma-Aldrich) 12% polyacrylamide (BioRad) gel in 1x Tris-Borate-EDTA running buffer (Sigma-Aldrich) at 200 V for 2.5 hours. Gels were stained with SYBR Safe (Invitrogen), diluted 1/10000 in running buffer, for 15 minutes under shaking and protected from light. Fragmented RNA smears were visualized by blue light transillumination and slices corresponding to the 60-120 nucleotide region were cut and placed into a 0.5 mL microcentrifuge tube (Eppendorf), in which a hole was punctured at the bottom. These tubes were placed into 2 mL microcentrifuge tubes (Eppendorf) and centrifuged at 17000 g for 1 min to crush slices into a slurry. RNA was extracted, precipitated and circularized as previously described^5^ with the addition of 10% v/v DMSO (Sigma-Aldrich) and 5% v/v PEG 8000 (New England Biolabs) during ligation.

### Reverse transcription and second strand synthesis

Ligation reactions were cleaned up with the Oligo Clean & Concentrator Kit and reverse transcribed using Superscript IV RT (Invitrogen) according to manufacturer’s instructions, using 0.5 µM of a gene-specific primer (sequences presented in **Supplementary Table 1**) bearing the M13 forward (−21) sequence followed by a unique molecular identifier (UMI, to distinguish unique molecules from PCR duplicates in the downstream analysis) of 10 random nucleotides as 5’ tag (5’-TGTAAAACGACGGCCAGTNNNNNNNNN), and extending the 55 ℃ incubation step to 1 hour.

Following RT, cDNAs were purified using 0.65 volumes of Ampure XP beads (Beckman) according to manufacturer’s instructions to select for large cDNAs containing several repeats. cDNAs were eluted off the beads using 20 µL of water and second strand synthesis reactions were assembled by adding 3 µL of a gene-specific primer (2 µM, sequences presented in **Supplementary Table 1**) with the T7 promoter sequence as 5’ tag (5’-TAATACGACTCACTATAGGG), 3 µL of 10x PfuUltra II Fusion buffer, 0.6 µL of PfuUltra II Fusion HS DNA polymerase (Agilent), 0.75 µL of dNTP (10 mM each, Roche) and 2.65 µL of nuclease-free water. Reactions were incubated at 95 ℃ for 5 min, 55 ℃ for 1 min, and 72 ℃ for 15 min, purified using 0.65 volumes of Ampure XP beads and eluted in 30 µL of water.

### PCR amplification and Illumina library preparation

Eluted dsDNA was amplified in three parallel PCR reactions using AmpliTaq Gold DNA polymerase (Applied Biosystems) with 5 µL of T7 promoter sequence primer (2 µM), 5 µL of M13 forward (−21) primer (2 µM), 5 µL of AmpliTaq MgCl_2_ solution, 5 µL of 10x AmpliTaq Gold buffer, 1 µL of AmpliTaq polymerase, 1.25 µL of dNTP (10 mM each) and 10 µL of purified dsDNA in a total reaction volume of 50 µL. Reactions were incubated at 95 ℃ for 5 minutes, followed by 30 cycles of 95 ℃ for 1 minute, 50 ℃ for 30 seconds, 72 ℃ for 2.5 minutes, and a final step at 72 ℃ for 6 minutes. PCR reactions were loaded on 1% agarose gels, the 300-500 bp region was excised and DNA was extracted using the MinElute Gel Extraction Kit (Qiagen).

DNA concentrations were measured using the Qubit dsDNA HS kit (Invitrogen) and 100 ng were used for paired-end Illumina sequencing library preparation using the KAPA HyperPlus Library Preparation Kit (Roche) and the KAPA Unique Dual-Indexed Adapater Kit (Roche), according to manufacturer’s instructions, starting directly with the End Repair and A-tailing step, replacing the double-sided selection step by a second Ampure Bead clean up step using 1 volume of beads, extending the 72 ℃ step during the PCR library amplification to 45 seconds, and performing the last clean up step with 0.7 volumes of beads. Concentrations of libraries were determined using the Qubit dsDNA HS kit, libraries were pooled equimolarly and sequenced using the Illumina V3 Miseq (2 x 300 cycles) or Illumina Nextseq 2000 (2 x 300 cycles) platform.

### Prediction of RdRp-trapping structures

Putative RdRp-trapping structures were predicted by a Python script using a sliding window approach and minimal free energy RNA structure prediction with the ViennaRNA^66^ package. A schematic representation of the method is shown in **Extended Data Fig. 1**. Nucleotides within an identically-sized window up- and downstream of the RdRp footprint, corresponding to the exiting and entering template strands respectively, were considered for sequence prediction. To mimic the close proximity of the entering 5’ and exiting 3’ template strand ends, the nucleotides in the RdRp footprint were replaced by an arbitrary sequence (AAGGGGGGAAAACCCCCCAA), constrained to fold as a stem-loop (constraint: `xx((((((xxxx))))))xx`, where `x`s denote nucleotides forbidden to pair, while open parentheses are forced to pair with the corresponding closed parentheses). Folding was then performed using the vrna_mfe function while disallowing lonely base pairs. After folding, the minimum free energy value was corrected by the minimum free energy value obtained with the arbitrary stem alone. The resulting standard Gibbs free energy change at 37°C 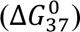 is denoted as ΔG. The script allows for variation in the size of the RdRp footprint (20 for figures in the main text), the position of the active site from the 5’ end of the sequence within the RdRp footprint (5 for figures in the main text) and the size of the flanking sequence window to consider. A 10-nucleotide window was used throughout the main text as, firstly, transient structures that would impact replication are expected to form close to the RdRp. Secondly, parts of the genome are bound by NP, which is only temporarily dislodged by the RdRp during replication. Prediction windows encompassing larger flanking regions have a higher risk of including NP-bound nucleotides, which would not be available to base pair. The interactive **Supplementary Fig. 1** shows the RNA structures predicted with a 10- or 15-nucleotide window for all HA sequences used. The script is available at https://github.com/dr-funk/trapped-RdRp/.

### Determining nature and frequency of insertions and deletions

Paired FASTQ files from Illumina paired-end sequencing were merged using AdapterRemoval^67^ (v. 2.3.2) and consensuses were extracted from repeat-containing reads and aligned to the reference by Bowtie2^68^ using the CirSeq software^69^, adapted to keep consensuses with insertions or deletions (indels) instead of discarding them. Due to low-frequency contamination of some transfections with A/Indonesia/5/2005 MBCS HA, leading to the presence of this sequence in some samples, including samples without RdRp, all consensuses bearing the A/Indonesia/5/2005 MBCS were excluded from further analysis (at most 0.04% of consensuses). Due to the size of the MBCS (24 nucleotides instead of 12) and the presence of a C in the A/Indonesia/5/05 MBCS stretch, these consensuses could be reliably separated from other consensuses bearing duplication-like insertions. SAM files containing the aligned consensuses were then processed using a Python script identifying consensuses with indels and calculating the coverage at each position (https://github.com/dr-funk/trapped-RdRp/). Consensuses containing more than four separate indel events or a one-nucleotide insertion and a one-nucleotide deletion occurring withing 5 nucleotides from each other where considered to result from contamination with a closely related HA template and discarded. To account for duplication of identical cDNAs by PCR, the UMI (10-nucleotide N stretch in the RT primer, see above) associated with each consensus was identified using the edlib alignment library^70^, and consensuses with the same UMI and alignment to the reference (as determined by alignment start position and CIGAR string) were filtered out to keep a single representative. Consensuses for which the UMI could not be identified were discarded.

Non-complex ambiguously positioned indels were identified and realigned by deducing every possible alignment for a given indel, calculating the stability of predicted RdRp-trapping structures at each possible position (on cRNA and vRNA, using the same approach as above), and realigning the indel to the position corresponding to the lowest free energy structure. For insertions, only alignments consistent with sequence duplication were kept. In HA templates with disrupted structures, if no structure is predicted in any possible position, the structures of the corresponding WT HA are used instead to facilitate comparison. Unambiguous indels were not realigned as, by definition, they have a single possible alignment. Complex indels were not realigned as they probably result from several independent indel events potentially occurring at several different positions. After prediction-based realignment, frequencies for each insertion were calculated based on the coverage at each given position. Interactive HTML files were generated using Plotly^71^ and jinja2 (https://palletsprojects.com/p/jinja). Numbers and figures in the text reflect insertion and deletion frequencies between positions 1041-1052 (or equivalent in H6) as they are the most relevant for MBCS acquisition.

### Analysis of cleavage site protein sequences

SAM files containing aligned consensuses were screened for insertions or deletions. Detection of contaminants and PCR duplicates was performed as above. If the difference between the number of inserted and deleted nucleotides was not a multiple of three, the cleavage site was considered to be frame-shifted. If the result was a multiple of three, the insertions and deletions were introduced into the corresponding reference sequence and the resulting sequence was translated to obtain the protein sequence. To determine whether insertions and deletions occurred in the HA cleavage site region, it was defined here as the translation of nucleotides between positions 1037 and 1057 for H5 references and 1032 and 1052 for H6 references. Frequencies were calculated by dividing counts by the maximum coverage observed in the RdRp-containing sample of each replicate and were not background-corrected. The interactive HTML file was generated using Plotly^71^ and jinja2 (https://palletsprojects.com/p/jinja).

### Unidirectional RdRp replication systems

Unidirectional RdRp-replication systems were established by RNP transfection as described above. To allow only vRNA to cRNA replication, a cRNA 3’ promoter deficient in the internal initiation step necessary for vRNA production (C1763A, cRNA sense)^72^ was used in combination with the A/Indonesia/5/2005 RdRp carrying a PA with the 51-72 loop replaced by a GGS linker (PA_Δ51-72_) ^73^ rendering the RdRp deficient in vRNA production. To allow only cRNA to vRNA replication, a vRNA 3’ promoter deficient in the terminal initiation step required for cRNA production (G2C and C9G, cRNA sense)^72^ was used in combination with the A/Indonesia/5/2005 RdRp carrying a PB1 priming loop deletion (PB1_Δ648-651_), which renders the RdRp deficient in vRNA production^74^. Due to unidirectional replication, different RNA senses had to be targeted for sequencing in RdRp-containing and non-RdRp (input) samples. In the vRNA to cRNA replication system, negative-sense RNA (vRNA template, PolI product) was targeted in the input samples, and positive-sense RNA (cRNA and mRNA, RdRp product) was targeted in the RdRp-containing samples. Opposite sense RNAs were targeted in the cRNA to vRNA replication system.

### Primer extension assay

Primer extension was performed as described previously^75^. In brief, primers specific for 18S rRNA (5’-GAGCCATTCGCAGTTTCACTGTAC), IN_RKKR_ vRNA (5’-CCTAGCACTGGCAATCATGATGG), or IN_RKKR_ cRNA/mRNA (5’-CATGATTGTGTCAACCTGCTCTG) were radioactively labelled using T4 polynucleotide kinase (New England Biolabs) and ɣ-^32^P-ATP (PerkinElmer). RT was performed on 2.5 µL of total RNA used for CirSeq processing (see above) using the radiolabeled primers and Superscript III reverse transcriptase (Invitrogen), resulting in labelled cDNAs of 96, 126 or 136 nucleotides for 18S rRNA, HA vRNA or HA cRNA respectively. HA mRNAs are amplified by the cRNA primer but run more slowly due to the presence of about 10-13 nucleotides added via cap-snatching alongside the 5’ cap ^76^. RT reactions were run on a denaturing urea polyacrylamide gel, which was imaged by overnight exposition of a BAS-IP MS 2040 E phosphor screen (Cytiva) scanned using a Typhoon FLA 9500 (GE Healthcare).

### Conservation of the 1043-1048 cRNA structure in H5 HPAIV precursors

For each H5 HPAIV outbreak with published HA sequences but no described LPAIV precursor, a putative precursor was identified by using the HPAIV HA as nucleotide BLAST (https://blast.ncbi.nlm.nih.gov/Blast.cgi) query against the nr/nt database and selecting the genetically closest HA sequence bearing a conserved RETR LPAIV cleavage site. For each putative precursor, the predicted cRNA RdRp-trapping structure at the position equivalent to 1045 in NL_RETR_ was determined using our transient structure prediction tool described above.

### Detection of putative non-homologuous recombination events

All consensuses containing complex insertions of nine or more nucleotides containing at least one C or U nucleotide (in cRNA sense) were identified. Sequences were aligned to the corresponding reference and selected as putative **non-homologuous recombination** events if the insert could not be explained by repeated duplication insertions. Putative **non-homologuous recombination** inserts were analysed by nucleotide BLAST (https://blast.ncbi.nlm.nih.gov/Blast.cgi) against the nr/nt database restricted to influenza A virus (NCBI:txid11320) or human (NCBI:txid9606) sequences to determine the putative origin of the insertion.

## Data availability

Next generation sequencing data that supports the findings of this study have been deposited in the European Nucleotide Archive as project PRJEB71774 (https://www.ebi.ac.uk/ena/browser/view/PRJEB71774).

Source data for Extended Data Fig. 4a are provided with the paper.

## Code availability

Python scripts used for data analysis are available at https://github.com/dr-funk/trapped-RdRp/.

## Supporting information

Suppelementary Table 2

## Acknowledgements

We thank M. Aguilar-Bretones, L. Bauer, B. Haagmans, S. Herfst, K. Schmitz and S. Thewessen for their critical reading and suggestions. This research was funded by ZonMw Off Road grant number 04510012010056, the NWO ENW XS grant OCENW.XS22.1.121, NWO ENW M1 grant OCENW.M.21.150, the European Union’s Horizon 2020 research and innovation program DELTA-FLU, grant agreement No. 727922, NIH/NIAID contract number HHSN272201400008C and 75N93021C00014, NID/NIAID grant DP2 AI75474, and NIH/NIAID R01 award 1R01AI177487. The funders had no role in study design, data collection and analysis, decision to publish or preparation of the manuscript.

## Author contributions

M.F. and M.R. conceived the project, M.F., M.I.S., A.J.W.tV. and M.R. designed experiments. M.I.S. generated the RNPs, M.F. extracted and processed the RNA, M.F and T.M.B. prepared the NGS libraries. M.F performed data analysis, M.F. wrote software with input from A.P.G. and A.J.W.tV. M.R. managed the project, M.F. and M.R. wrote the initial manuscript, M.F., M.I.S., T.M.B., A.C.M.dB., A.P.G., R.A.M., A.J.W.tV. and M.R. edited the manuscript. Funding was acquired by M.F., A.J.W.tV. and M.R.

## Declaration of interests

The authors declare no competing interests.

## Additional information

Supplementary Information is available for this paper.

Correspondence and requests for materials should be addressed to Mathis Funk or Mathilde Richard.

## Extended Data

**Extended Data Figure 1:**
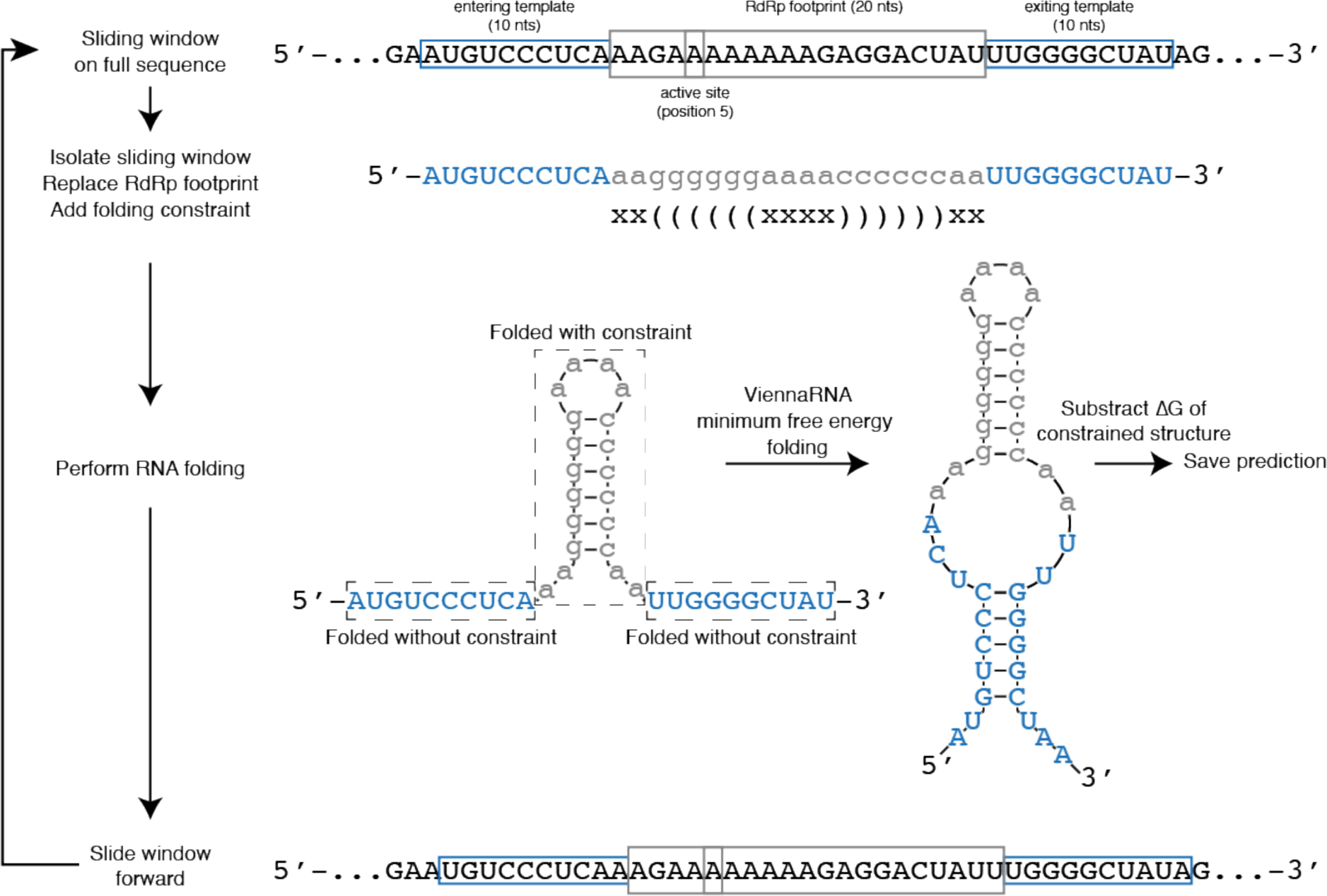
Predicting transient RdRp-trapping RNA structures. Schematic representation of the transient structure prediction method. A sliding window containing the RdRp footprint (20 nts here) as well as a set length of RNA on either side (10 here) are isolated and the sequence within the RdRp footprint replaced by an arbitratry sequence. RNA folding is performed with a folding constraint on the arbitrary sequence, x indicates a nucleotide that is not allowed to pair while corresponding parentheses indicate nucleotides forced to pair together. The constrained structure brings the 3’ end of the exiting template into close proximity of the 5’ end of the entering template (as would occur during replication). After folding, the free energy change (ΔG) of the transient structure is corrected by subtracting the ΔG of the constrained sequence. The window then slides forward by one nucleotide and the process is repeated.

**Extended Data Figure 2:**
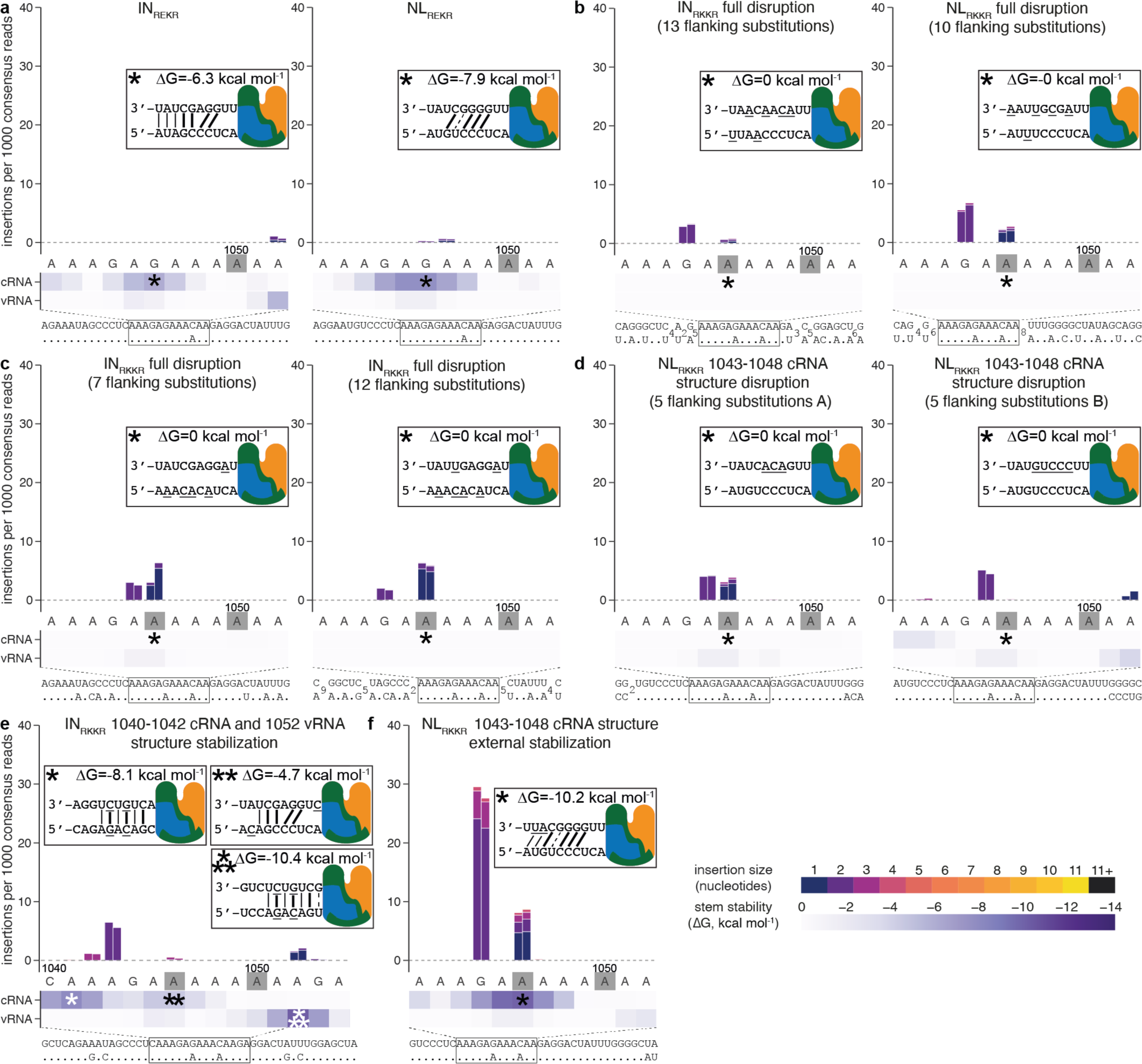
Insertion frequencies observed with additional IN and NL mutants. **a-f** Insertion frequencies observed in IN_REKR_ (left) or NL_REKR_ (right) (**a**), in an RKKR cleavage site and substitutions disrupting all predicted RdRp-trapping structures in IN_RKKR_ (left) and NL_RKKR_ (right) (**b**), in IN_RKKR_ with substitutions disrupting all predicted RdRp-trapping structures (**c**), in NL_RKKR_ with substitutions disrupting the 1043-1048 cRNA structure (**d**), in IN_RKKR_ with substitutions stabilizing the 1040-1042 cRNA/1052 vRNA structures (**e**), or in NL_RKKR_ with substitutions stabilizing the 1043-1048 cRNA structure by creating additional basepairs at the edge most distant of the RdRp (**f**). Using the same representation as **Fig. 2**. Due to size constraints, some alignments were broken into chunks, each separated by the indicated number of identical positions.

**Extended Data Figure 3:**
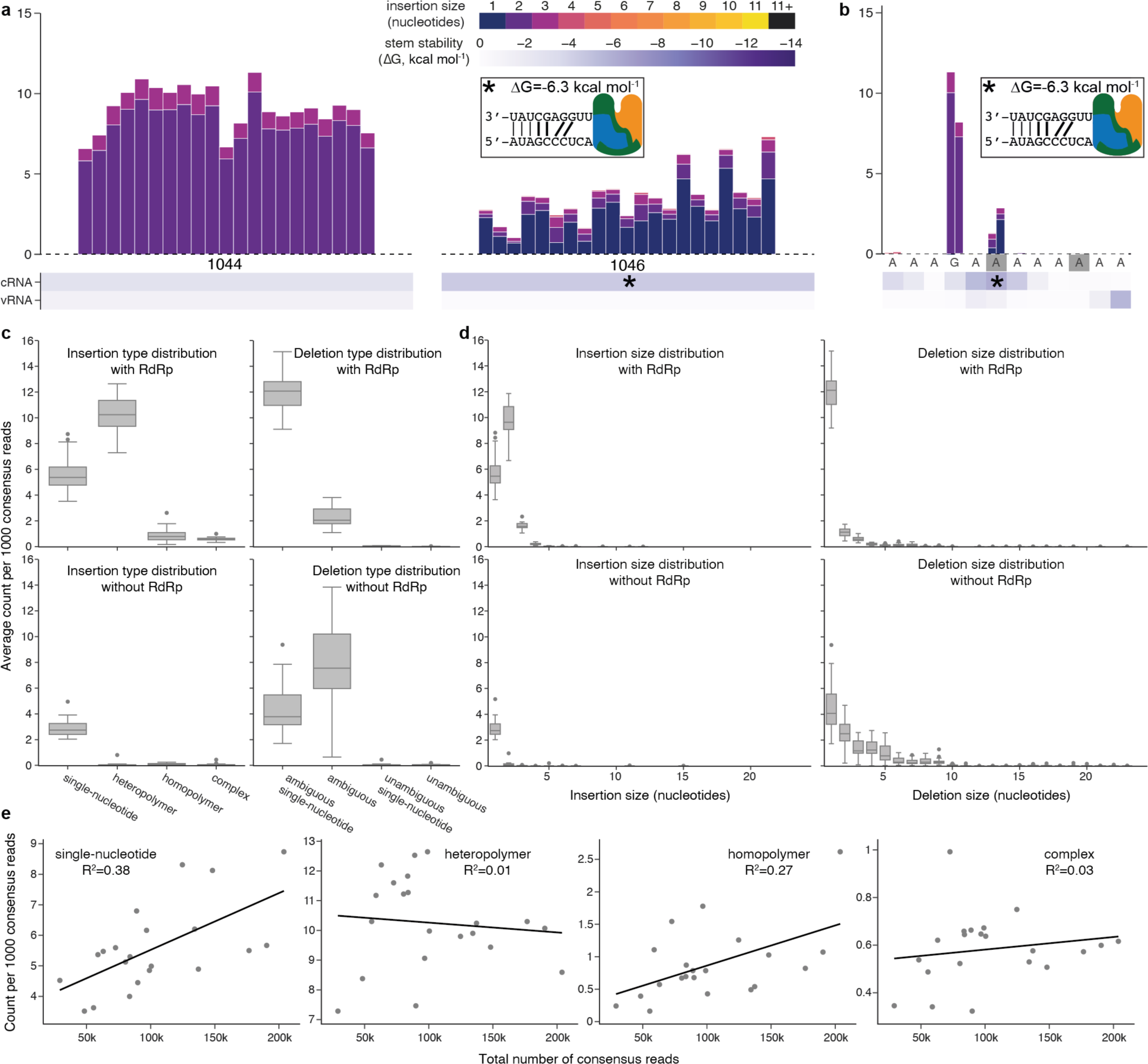
Insertion frequencies are consistent across replicates and do not correlate with number of consensuses. **a-b,** Insertions frequencies observed in IN_RKKR_ in 21 independent biological replicates (**a**) or using different primers for RNA processing (**b**). Using the same representation as **Fig. 2**, scales at the top. **c-d,** Uncorrected insertion frequency distribution by type of insertion (**c**) or size of insertion (**d**) for all 21 IN_RKKR_ replicates. The upper and lower bounds of the box show the 3^rd^and 1^st^ quartile respectively (linear method) while the central bar shows the median. Whiskers show the highest or lowest value within a 1.5x interquartile range from the 3^rd^ and 1^st^ quartile respectively. Outliers beyond that range are shown as points. **e,** Scatter plots showing insertion frequencies as a function of total number of consensus reads for each replicate. Insertions were split by type, and linear regression lines were plotted, with the value of the correlation coefficient R^2^ indicated in each panel.

**Extended Data Figure 4:**
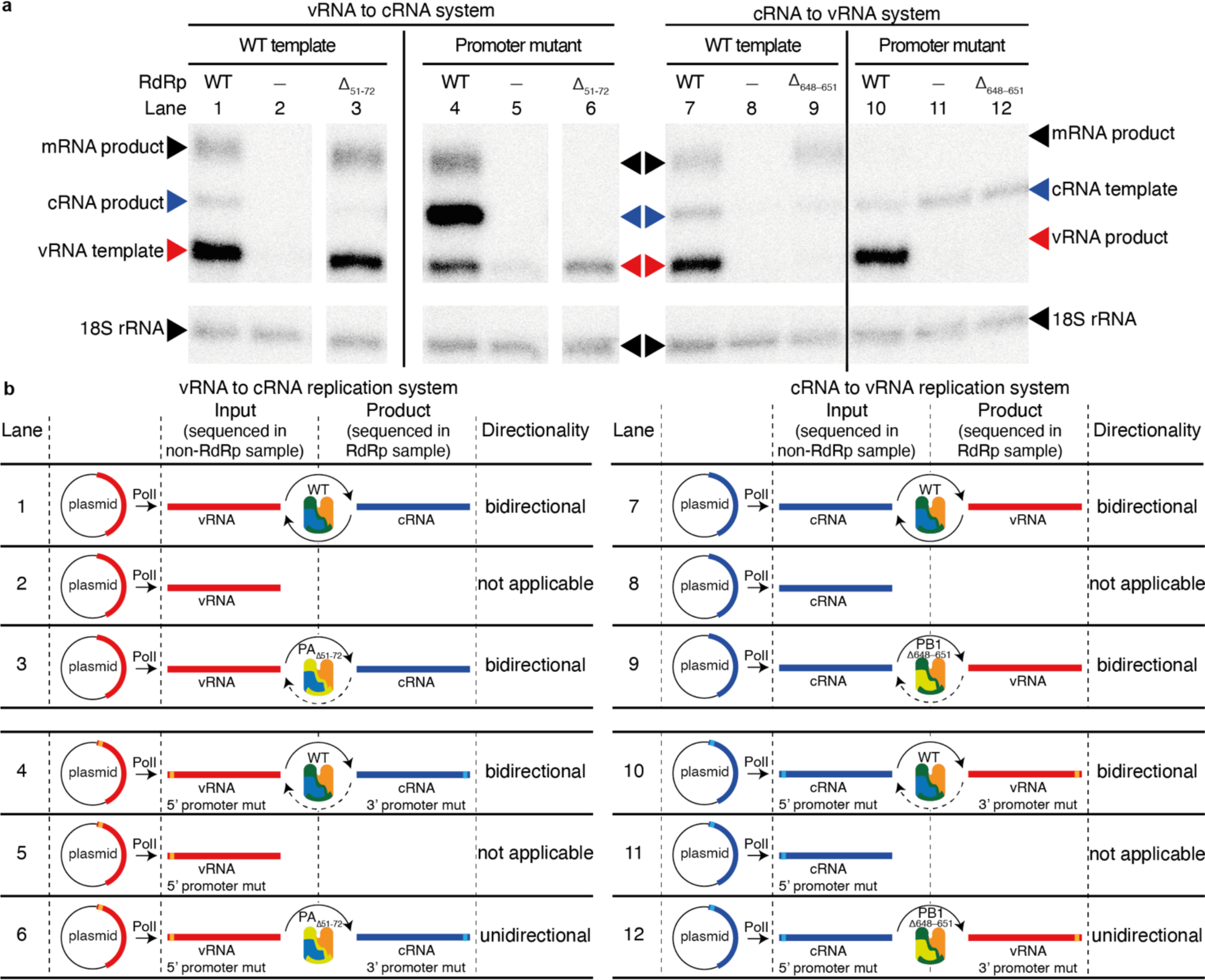
A unidirectional RNP-based influenza replication system. **a,** Primer extension assay discriminating influenza virus RNA species produced with combinations of mutant or WT RNA and RdRp. Expected height for each cDNA product are indicated by arrowheads, with the cRNA and vRNA arrowheads colored as in **b**. 18S rRNA was used as a loading control. **b,** Schematic overviews of the conditions used for the unidirectional assay, including the observed directionality for each combination of mutant or WT RNA template and RdRp. Lane numbers correspond to lanes in **a**.

**Extended Data Table 1:**
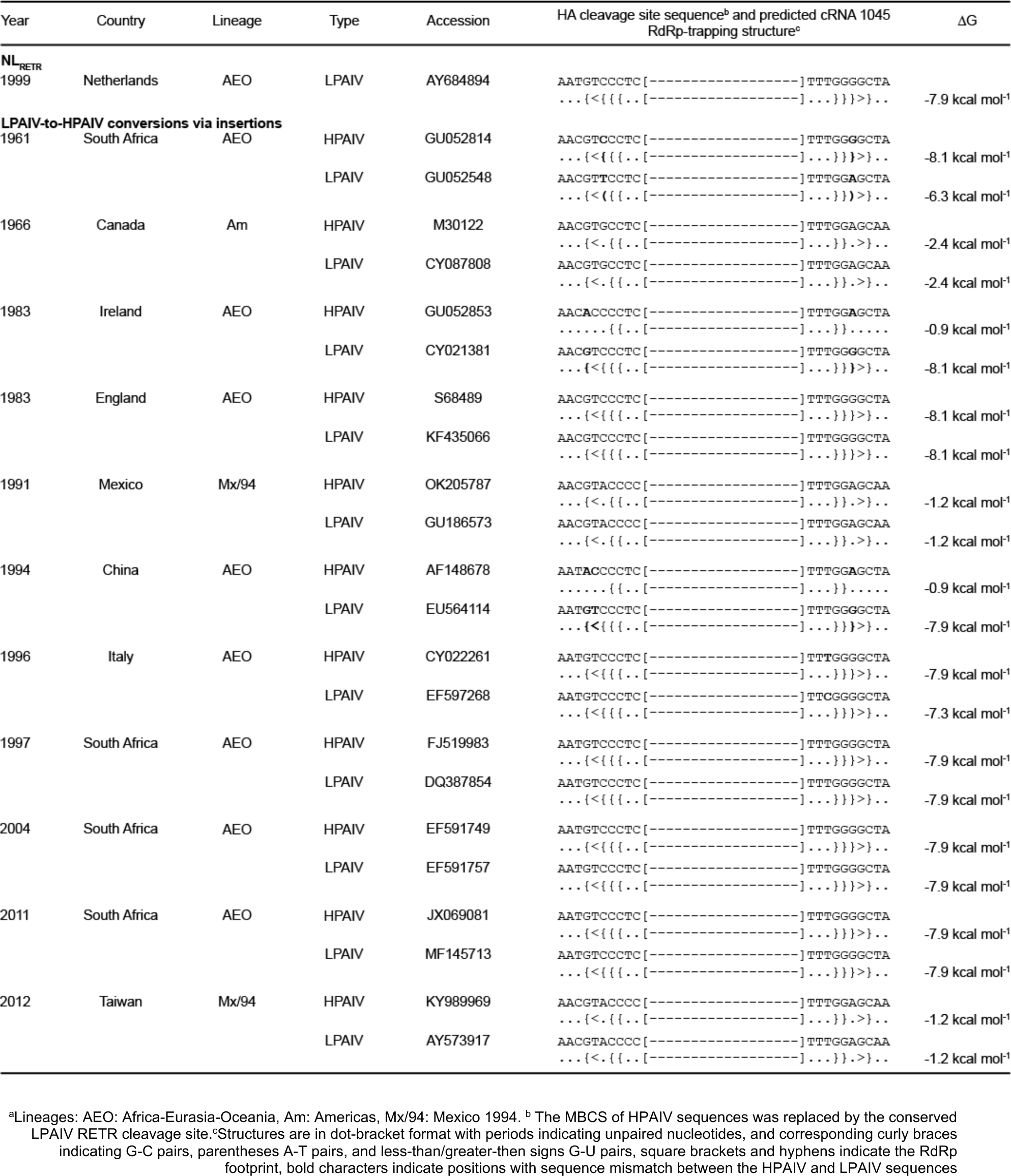
Predicted 1045 cRNA RdRp-trapping structures of NL_RETR_ and putative H5 precursors.

## Supplementary information

**Supplementary Notes 1-10**

Additional text, including discussion of potential non-homologous recombination events, the presence of the 1043-1048 cRNA structure in HPAIV and the impact of sequence and structure on deletions.

**Supplementary Table 1**

**Primers used for RT and second-strand synthesis.** In order of appearance in the main text. For IN_RKKR_, primers are sorted by RNA sense targeted for sequencing. All primer sequences are given in 5’ to 3’ sense.

**Supplementary Table 2**

**Putative non-homologous recombination events.** This table shows the sequences inserted during putative non-homologous recombination events as well as their most likely origin via BLAST.

**Supplementary Figure 1**

**Predicted RdRp-trapping structures in all HA templates**. Interactive heatmaps showing the predicted RdRp-trapping structures for all templates used in this study. For non-RETR/IETR templates, only the structures formed in the region surrounding the HA cleavage site are shown.

**Supplementary Figure 2**

**Interactive insertion frequency graphs for all HA templates**. Interactive graphs showing the uncorrected insertion frequencies observed in each HA template used in this study. They offer additional information when compared to the graphs of the main text such as coverage, a detailed breakdown of insertion type and alignment of insertions.

**Supplementary Figure 3**

**Effects of insertions on HA cleavage site protein sequence**. Interactive stacked bar graphs showing resulting protein sequences at the HA cleavage site for each sample. Protein sequences are grouped into WT (blue), in-frame insertion/deletion (red), and out-of-frame insertion/deletion (grey). Hovering over a bar displays the corresponding frequency and protein sequence.

**Supplementary Figure 4**

**Interactive deletion frequency graphs for all HA templates**. Interactive graphs showing the uncorrected deletion frequencies obtained with all templates used in this study. They offer additional information when compared to the graphs of the main text such as coverage, a detailed breakdown of deletion type and alignment of deletions.

**Supplementary Data 1**

**Sequences of all HA templates.** This file contains all full segment sequences of the templates used for this study in FASTA format

**Supplementary Data 2**

**Insertion and deletion frequencies by type**. This tab-separated text file contains the uncorrected insertion and deletion frequencies for each replicate and each HA template. The frequencies are sorted into the insertion and deletion types described in the text.

**Supplementary Video**

**The trapped-RdRp model**. A detailed animation showing how the most commonly observed insertion, 1044GA (cRNA sense), could be explained by the trapped-RdRp model. Using the same representation as **Fig. 1**, RNA structures at the bottom are in dot-bracket format, `{}` for GC, () for AU, `<>` for GU base pairs.

## Notes

https://www.ebi.ac.uk/ena/browser/view/PRJEB71774

https://github.com/dr-funk/trapped-RdRp

### Competing Interest Statement

The authors have declared no competing interest.

