## Supplementary Figure 1 for "Transient RNA structures underlie highly pathogenic avian influenza virus genesis": supplementary_figure_1-predictions.html

×


  

parameters

IN_X_ main text

IN_RETR_
  


IN_RKKR_
  


IN_RKKR_ 1043-1048 cRNA stem destabilization
  


IN_RKKR_ with NL_RETR_ 1043-1048 cRNA stem
  

IN_X_ Extended Data

IN_REKR_
  


IN_RKKR_ full destabilization (13 substitutions)
  


IN_RKKR_ destabilization (7 substitutions)
  


IN_RKKR_ full destabilization (12 substitutions)
  


IN_RKKR_ Gs/Gd-specific stem stabilization
  

NL_X_ main text

NL_RETR_
  


NL_RKKR_
  


NL_RKKR_ 1043-1048 cRNA stem destabilization
  


NL_RKKR_ 1043-1048 cRNA stem internal stabilization
  

NL_X_ Extended Data

NL_REKR_
  


NL_RKKR_ full destabilization (10 substitutions)
  


NL_RKKR_ 1043-1048 cRNA stem destabilization (5 substitutions A)
  


NL_RKKR_ 1043-1048 cRNA stem destabilization (5 substitutions B)
  


NL_RKKR_ 1043-1048 cRNA stem external stabilization
  

SW_X_ main text

SW_IETR_
  


SW_RKKR_
  


SW_RKKR_ 1043-1048 cRNA stem destabilization
  


SW_RKKR_ with NL_RETR_ 1043-1048 cRNA stem

☰ Go To...

### Predicted RdRp-trapping structures for all constructs of Funk et al. 2024

_made using slidingfold version 23/20/09_

This file contains transient RdRp-trapping RNA structure predictions for all sequences used in Funk et al 2024, it might need a few seconds to finish loading. While loading the sidebar menu will not work. Similarly, the sidebar makes use of jQuery and will not be fully functional when offline.

The following parameter models were used for folding (from inner to outer ∆G heatmap):

| parameter set | footprint | active site location | nts on each side |
| --- | --- | --- | --- |
| 1 | 20 | 5 | 10 |
| 2 | 20 | 5 | 15 |

Lonely RNA pairs were forbidden and the RdRp footprint was replaced by an arbitrary sequence with folding constraint (x is unpaired, parentheses indicated forced pairs):  

AAGGGGGGAAAACCCCCCAA  
 xx((((((xxxx))))))xx

The "Go To..." menu at the top left allows to quickly navigate to a specific graph via a sidebar table of content. Checkboxes next to each graph allow to show or hide the graph and the buttons at the top allow to hide or show all graphs at once. Clicking on the name of a hidden graph will set it to be shown and move the view over the graph. Clicking on the name of a subcategory will cause all its graphs to be shown and move the view to the first graph of the subcategory. To close the sidebar, simply click anywhere outside it.

Hovering over different parts of the graphs provides additional information, such as model parameters, ∆G, and predicted RdRp-trapping structure. To zoom into a particular part of the figure, click-and-drag your mouse across is. To reset the zoom, double-click in an empty area of the graph. To see the entire graph, double-click in an empty area while at the default zoom level. Zoom levels can also be controlled via the menu in the upper-right corner of the graph.

At the bottom of the graph, the cursor changes into a horizontal double arrow, clicking and dragging allows to move the displayed region left or right.

xml version="1.0" encoding="utf-8"?


footprint: 20 nts, active site in position 5
10 nts on each side for folding
position 1045 ∆G= -7.90 kcal mol
-1
AUGUCCCUCAAAGAGAAACAAGAGGACUAUUUGGGGCUAU
..{<{{{...[---GAAACAAGAG-----]..}}}>}...

Predicted cRNA structures(10 nucleotide window)
Predicted cRNA structures(15 nucleotide window)
Predicted vRNA structures(15 nucleotide window)
Predicted vRNA structures(10 nucleotide window)
Amino acid sequence
cRNA sequence
cRNA position
Structure infobox appears when hovering over heatmap
Nucleotides:
Amino acids (ClustalX color scheme):
RNA structures ∆G (kcal mol
-1
)
Adenine
Purines
Hydrophobic
Uracyl
Cytosine
Guanine
Alanine
Leucine
Isoleucine
Phenylalanine
Methionine
Lysine
Arginine
Glutamate
Aspartate
less stable
more stable
Tryptophane
Valine
Asparagine
Threonine
Serine
Glutamine
Cysteine
Histidine
Tyrosine
Stop codon
Proline
Glycine
cRNA position
vRNA sequence


RdRp model parameters
Position and free energy,lower is more stable
Sliding window size


RdRp footprint


Nucleotide in RdRp active site

Matched bracketsrepresent base pairs:

Periods representunpaired nucleotides


Pyrimidines


Polar


Aromatic


Positivecharge


Negativecharge
0
-14
-12
-10
-8
-6
-4
-2


990

1000

1040

1050

1040


990

1000

1040

1050

1040


C

C

U

C

A

A

A

G

A

G

A

A

A

C

A

A

G

A

G

G

A

C

U

A

P

Q

R

E

T

R

G

L

G

G

A

G

U

U

U

C

U

C

U

U

U

G

U

U

C

U

C

C

U

G

A

U
1060
1060


{}
()
<>

G-C base pair
A-U base pair
G-U base pair

Strength of
base pair


C
U
A
G


S


L


F


Y


W


C


\*


P


Q


R


H


I


K


N


T


V


E


D


A


G


M

Full-length predictions are provided for all LPAIV HAs, with a window of 81 nucleotides encompassing the HA cleavage site being shown by default. For mutants, the 5'- and 3'-most positions showing an altered stem were identified and only positions in the corresponding region and/or the 81 nucleotide default window are shown.
