## Supplementary Figure 2 for "Transient RNA structures underlie highly pathogenic avian influenza virus genesis": supplementary_figure_2-insertions.html

×


About

show explanatory text

IN_X_ main text

IN_RETR_
  


IN_RKKR_
  


IN_RKKR_ 1043-1048 cRNA stem destabilization
  


IN_RKKR_ with NL_RETR_ 1043-1048 cRNA stem
  

IN_X_ Extended Data

IN_REKR_
  


IN_RKKR_ second primer set
  


IN_RKKR_ full destabilization (13 substitutions)
  


IN_RKKR_ destabilization (7 substitutions)
  


IN_RKKR_ full destabilization (12 substitutions)
  


IN_RKKR_ Gs/Gd-specific stem stabilization
  


IN_RKKR_ 21 independent repeats
  

NL_X_ main text

NL_RETR_
  


NL_RKKR_
  


NL_RKKR_ 1043-1048 cRNA stem destabilization
  


NL_RKKR_ 1043-1048 cRNA stem internal stabilization
  

NL_X_ Extended Data

NL_REKR_
  


NL_RKKR_ full destabilization (10 substitutions)
  


NL_RKKR_ 1043-1048 cRNA stem destabilization (5 substitutions A)
  


NL_RKKR_ 1043-1048 cRNA stem destabilization (5 substitutions B)
  


NL_RKKR_ 1043-1048 cRNA stem external stabilization
  

unidirectional main text

IN_RKKR_ cRNA to vRNA WT RdRp
  


IN_RKKR_ cRNA to vRNA promoter mutant PB1_∆648-651_ mutant RdRp
  


IN_RKKR_ vRNA to cRNA WT RdRp
  


IN_RKKR_ vRNA to cRNA promoter mutant PA_∆51-72_ mutant RdRp
  

SW_X_ main text

SW_IETR_
  


SW_RKKR_
  


SW_RKKR_ 1043-1048 cRNA stem destabilization
  


SW_RKKR_ with NL_RETR_ 1043-1048 cRNA stem

☰ Go To...

### Interactive insertion frequency graphs

This file contains interactive versions of all insertion graphs of Funk et al 2024, it might need a few seconds to finish loading. While loading the sidebar menu will not work. Similarly, the sidebar makes use of jQuery and will not be fully functional when offline.

The "Go To..." menu at the top left allows to quickly navigate to a specific graph via a sidebar table of content. Checkboxes next to each graph allow to show or hide the graph and the buttons at the top allow to hide or show all graphs at once. Clicking on the name of a hidden graph will set it to be shown and move the view over the graph. Clicking on the name of a subcategory will cause all its graphs to be shown and move the view to the first graph of the subcategory. To close the sidebar, simply click anywhere outside it.

To zoom into a particular part of the figure, click-and-drag your mouse across is. To reset the zoom, double-click in an empty area of the graph. To see the entire graph, double-click in an empty area while at the default zoom level. Zoom levels can also be controlled via the menu in the upper-right corner of the graph.

At the bottom of the graph, the cursor changes into a horizontal double arrow, clicking and dragging allows to move the displayed region left or right.

The graphs default to showing the HA cleavage site and extend on either side until coverage drops to less than 1% of the respective maximum for all replicates and samples of the graph. While there are some insertions with high apparent frequencies in the flanking regions, these frequencies are likely to be inflated due to the low coverage in these regions, since insertion frequency is calculated with regards to the actual coverage at the same position.

xml version="1.0" encoding="utf-8"?


replicate 2, size 3 nts
1.69/1000 consensus reads
aaaga---aaaaa: complex
aaagaGAAaaaaa 0.92‰
aaaga---aaaaa: homopolymer
aaagaAAAaaaaa 0.76‰
aaaga---aaaaa: complex
aaagaGAGaaaaa 0.01‰

Insertion frequencies in
RdRp-containing samples
are plotted as positive
Legend contains only insertion sizes observed 
for this template
Click on a legend item to toggle display of 
corresponding data
Double-click on a shown legend item to show only 
corresponding data, double-click on a hidden legend 
item to show all data 

Replicate 1

Replicate 2
Coverage RdRp-containing samples
is plotted as positive


Coverage background/input samples
is plotted as negative

Insertion frequencies in background
samples are plotted as negative
RNA sequence with substitutions
compared to LPAIV highlighted
Predicted RdRp-trapping
RNA structure heatmap


Insertion infobox
 appears when hovering over a bar

Structure infobox
 appears when hovering over heatmap

Legend
 on the right might be scrollable if too long

Total frequency of insertions of
this size at this position


Stability of RNA structure
(lower is more stable)

{}
()
<>
.

G-C base pair
A-U base pair
G-U base pair
unpaired nucleotide

Strength of
base pair

RdRp footprint

Reference

Inserted sequence


Insertion type


Nucleotide in RdRp active site


Frequency of this insertion

Breakdown of all insertions of
this size at this position, with
one alignment per insertion

insertion size
1 nts insertions
2 nts insertions
3 nts insertions
4 nts insertions
5 nts insertions
6 nts insertions
9 nts insertions
11 nts insertions
15 nts insertions
coverage replicate 1
coverage replicate 2


cRNA position 1046 dG= -7.90 kcal mol
-1
AUGUCCCUCAAAGAAAAAAAAGAGGACUAUUUGGGGCUAU
..{<{{{...[---AAAAAAAGAG-----]..}}}>}...


Amino acid sequence 


0


10


20


30


40


A


A


A


G


A


A


A


A


A


A


A


A


cRNA


vRNA


−50k


0


50k


100k


insertions per 1000 consensus reads
coverage


Q


 


 


R


 


 


K


 


 


 


 

RNA structure predictions were made using ViennaRNA following the same method as the Python script available at https://github.com/dr-funk/slidingfold, with an RdRp footprint of 20 nucleotides with the active site in position 5, and a window size of 10 nucleotides on each side of the RdRp.
