## Supplementary Figure 3 for "Transient RNA structures underlie highly pathogenic avian influenza virus genesis": supplementary_figure_3-protein.html

×


About

show explanatory text

IN_X_ main text

IN_RETR_
  


IN_RKKR_
  


IN_RKKR_ 1043-1048 cRNA stem destabilization
  


IN_RKKR_ with NL_RETR_ 1043-1048 cRNA stem
  

IN_X_ Extended Data

IN_REKR_
  


IN_RKKR_ second primer set
  


IN_RKKR_ full destabilization (13 substitutions)
  


IN_RKKR_ destabilization (7 substitutions)
  


IN_RKKR_ full destabilization (12 substitutions)
  


IN_RKKR_ Gs/Gd-specific stem stabilization
  

NL_X_ main text

NL_RETR_
  


NL_RKKR_
  


NL_RKKR_ 1043-1048 cRNA stem destabilization
  


NL_RKKR_ 1043-1048 cRNA stem internal stabilization
  

NL_X_ Extended Data

NL_REKR_
  


NL_RKKR_ full destabilization (10 substitutions)
  


NL_RKKR_ 1043-1048 cRNA stem destabilization (5 substitutions A)
  


NL_RKKR_ 1043-1048 cRNA stem destabilization (5 substitutions B)
  


NL_RKKR_ 1043-1048 cRNA stem external stabilization
  

unidirectional main text

IN_RKKR_ cRNA to vRNA WT RdRp
  


IN_RKKR_ cRNA to vRNA promoter mutant PB1_∆648-651_ mutant RdRp
  


IN_RKKR_ vRNA to cRNA WT RdRp
  


IN_RKKR_ vRNA to cRNA promoter mutant PA_∆51-72_ mutant RdRp
  

SW_X_ main text

SW_IETR_
  


SW_RKKR_
  


SW_RKKR_ 1043-1048 cRNA stem destabilization
  


SW_RKKR_ with NL_RETR_ 1043-1048 cRNA stem

☰ Go To...

### Amino acid HA cleavage site sequences

This file contains interactive graphs showing the amino acid cleavage site sequences resulting from the insertions and deletions observed for each sample.

The "Go To..." menu at the top left allows to quickly navigate to a specific graph via a sidebar table of content. Checkboxes next to each graph allow to show or hide the graph and the buttons at the top allow to hide or show all graphs at once. Clicking on the name of a hidden graph will set it to be shown and move the view over the graph. Clicking on the name of a subcategory will cause all its graphs to be shown and move the view to the first graph of the subcategory. To close the sidebar, simply click anywhere outside it.

To zoom into a particular part of the figure, click-and-drag your mouse across is. To reset the zoom, double-click in an empty area of the graph. To see the entire graph, double-click in an empty area while at the default zoom level. Zoom levels can also be controlled via the menu in the upper-right corner of the graph.

At the bottom of the graph, the cursor changes into a horizontal double arrow, clicking and dragging allows to move the displayed region left or right.

xml version="1.0" encoding="utf-8"?


0

200

400

600

800

1000


2

1

NL
RKKR
 

 

replicate

Frequency of consensuses with wild-type HA cleavage site sequence
Frequency of consensuses with in-frame indelsin the HA cleavage site
Frequency of consensuses with out-of-frame indelsin the HA cleavage site


Infobox appears when hovering over a bar:
Sample andreplicate number
Resulting amino acid cleavage site sequence
Size of indel: Absent if wild-type In amino acids (aa) if in-frame In nucleotides (nts) if out-of-frame
Frequency of this amino acid cleavage site sequence


NLinsertion (+1 aa) (PQRRKKRG)3.96 per 1000 consensuses
RKKR
(1)
