## Supplementary Notes for "Transient RNA structures underlie highly pathogenic avian influenza virus genesis"

### Table of Contents


**SN1:** Quantifying rare insertions at the HA cleavage site 1

**SN2:** Non-homologous recombination events 2

**SN3:** Insertion frequencies are comparable across 21 replicates 4

**SN4:** Background insertions are affected by sequence but not predicted RNA structures 5

**SN5:** Inconsistent single-nucleotide and homopolymer insertions in disruption mutants 6

**SN6:** Differences in insertion frequencies depending on amplified RNA sense 7

**SN7:** H5 HPAIV precursors contain the 1043-1048 cRNA structure 8

**SN8:** Deletions are promoted by A/U stretches 9

**SN9:** Deletion frequencies are not affected by RNA structures 10

**SN10:** Deletions by the RdRp occur mostly on cRNA 10

**Supplementary Table 1**: Primer sequences used in this study 11


#### Supplementary Note1: Quantifying rare insertions at the HA cleavage site

RNPs were generated in two biological replicates using the A/Indonesia/5/2005 RdRp (PB2, PB1, PA) and NP, and an HA vRNA as template. In parallel, background insertion levels were quantified in two biological replicates by omitting the plasmids coding for the RdRp during transfection. Background insertions are expected to be primarily due to errors by the human DNA polymerase I (PolI) during generation of the initial RNA templates from plasmid DNA. In order to reliably identify rare insertions, we used a custom circular sequencing method[5]. RNA molecules are circularized, leading to cDNA tandem repeats upon reverse transcription. After sequencing, repeats are collapsed into consensuses and only consensus mutations are kept, as those are most likely present in the original RNA molecule.

Insertions in repetitive sequences can often be aligned to the reference in several valid ways. Aligning these insertions consistently requires an arbitrary rule for standardizing insertions placement. For potential duplication events, i.e. the inserted sequence is identical to neighboring sequences, we identified all possible RdRp-trapping structures, in both cRNA and vRNA sense, that could lead to a duplication according to our model (see **Methods**). Each insertion was then aligned to the position of the most stable predicted RNA structure. In addition to standardizing insertion placement, this allowed us to test compatibility with our model by identifying putative RdRp-trapping structures that might be involved in insertions. Insertion frequencies were then corrected by subtracting the background frequencies observed without RdRp, with insertions realigned the same way, to allow direct comparison between repeats. All raw non-corrected insertion frequencies for RdRp-containing and background samples are provided as an interactive figure in **Supplementary Fig. 2**, alongside additional information such as alignments, predicted structures, and coverage at each position.

**Supplementary Note 2: Non-homologous recombination events**

To date, no confirmed non-homologous recombination (NHR) events have been detected in H5 or H6 HAs in nature, though a twelve-nucleotide insertion in the H5 A/tern/South Africa/1961 HPAIV HA (between positions equivalent to 1050-1051 in H5 LPAIV HAs) is suspected to result from NHR due to low similarity to adjacent sequences[2]. While almost all insertions observed at the H5 and H6 cleavage sites were clear duplications, characterized by a sequence composition similar to the surrounding sequences, eleven large (defined here as nine or more nucleotides) insertions of low similarity were detected (**Supplementary Table 2**), suggesting they may have resulted from NHR. These insertions occurred without deletions and most had no or only one ambiguously placed nucleotide at the edges. Similar insertions without deletion and with low ambiguity were observed in the known NHR events in H7 HPAIVs[12-16]. While we cannot exclude artifactual NHR-like consensuses resulting from artificial random ligation of randomly fragmented RNA molecules, these would be expected to present artefactual deletions or duplications in addition of the low-similarity insertion, resulting from ligating differently fragmented HA cleavage site molecules to either side of an exogenous RNA molecule.

Of the eleven putative NHR events, eight occurred in $NL_{RKKR}$-, two in $IN_{RKKR}$- and one in $SW_{RKKR}$-based samples. Putative NHR insertions were observed at a range of positions, with seven of eleven insertions occurring in the eight-nucleotide A stretch of the RKKR cleavage site (H5 positions 1045-1052). Of the other four, three occurred upstream, and one downstream of the RKKR A stretch. Eight putative NHR insertions were in frame, but four of these contained a stop codon in the insert. All but two inserted sequences showed identity with at least one human genome sequence via BLAST, though, due to their limited sizes (12-24 nucleotides), it is difficult to determine their origin with certainty. Six out of eleven insertions showed identity with ribosomal RNA (rRNA) sequences and two out of eleven with transfer RNA (tRNA) sequences, both of which have previously been suggested to be the origin of NHR-derived insertions in H7 HA genes[14-16] (**Supplementary Table 2**). Strikingly, two insertions were observed twice. Firstly, the exact same unambiguous tRNA-derived insertion was found at neighboring positions (1046-1047 and 1047-1048) in two samples with different $IN_{RKKR}$-based templates. Secondly, two overlapping 18S rRNA-derived insertions (nucleotides 482-463 and 483-464 of the 18S rRNA) were inserted in two different samples with different $NL_{RKKR}$-based templates, again at neighboring positions (1042-1043 and 1044-1045). These repeated insertions of (near-)identical sequences in adjacent locations fit with the observation of (near-)identical NHR events in distinct H7 influenza viruses in nature[2] and the hypothesis that influenza NHR may be guided by sequence-specific interaction with snoRNAs[16].

Intriguingly, six of eleven potential NHR insertions were observed in samples without RdRp, suggesting that recombination in influenza virus genomes might not be mediated by the viral RdRp, but might occur non-replicatively, for example following RNA strand breakage in cells. Non-replicative NHR has been previously shown to occur in pestiviruses[48-50] and poliovirus[51] and proposed to occur via strand breakage and rejoining.

When analyzed in cRNA sense, all putative NHR insertions corresponded to the reverse-complement of cellular RNAs, in accordance to what has been observed for NHR in H7 HAs[16], suggesting that recombination in H5 HAs also occurs in a vRNP (non-replicative NHR) or vRNA to cRNA replication (replicative NHR) context.

**Supplementary Note 3: Insertion frequencies are comparable across 21 replicates**

To evaluate the variation between independent experiments, $IN_{RKKR}$ was included as internal control in every round of RNP generation and circular sequencing, resulting in 21 independent biological replicates (**Extended Data Fig. 3a).** The two replicates with the most consensus reads being used as representatives in the main text. Uncorrected insertion frequencies between RdRp-containing replicates were consistent across the 21 replicates, with little variation for heteropolymer insertions, yet more variation for single-nucleotide and especially for homopolymer insertions (**Extended Data Fig. 3c, top left panel**). We also observed little variation for heteropolymer insertion frequencies when using a second set of primer sequences for RT and second strand synthesis (8.5-10.1‰ to 8.3-11.4‰), while single-nucleotide (2.4-5.1‰ to 0.2-2.1‰) and homopolymer insertions (1.0-2.5‰ to 0.4-0.6‰) were more variable (**Extended Data Fig. 3b**).The size distribution of insertions was consistent among the 21 replicates, with most insertions in RdRp-containing samples being 2 nucleotides long and insertion frequencies rapidly declining as size increases (**Extended Data Fig. 3d, top left panel**). The number of consensus reads obtained for RdRp-containing samples ranged from about 30,000 to 200,000, but no correlation between consensus read numbers and insertion frequencies was observed (**Extended Data Fig. 3e**).

**Supplementary Note 4: Background insertions are affected by sequence but not predicted RNA structures**

Background insertion frequencies observed in the samples without RdRp increased significantly when the RETR was replaced with the RKKR cleavage site (**Supplementary Fig. 2**). While in RdRp-containing samples, this change led to a higher increase in insertion frequencies in NL than in IN templates (**Fig. 2a,b**), no differences between IN (0.0-0.0‰ to 3.3-3.8‰) and NL templates (0.0-0.1‰ to 3.8-4.6‰) were observed in non-RdRp samples. Background insertions were almost exclusively insertions of 1 or 2 As in the 1045-1052 A stretch ($IN_{RKKR}$: 3.3-3.7‰, $NL_{RKKR}$: 3.7-4.5‰), with virtually no larger homopolymer (<0.0‰) or heteropolymer insertions (<0.1‰). This observation was confirmed in the 21 $IN_{RKKR}$ replicates (**Extended Data Fig. 3c-d, bottom left panels).** This is consistent with PolI errors as described previously[44], suggesting that background insertions are indeed introduced by PolI.

Importantly, while insertion frequencies in RdRp-containing samples varied when predicted RdRp-trapping RNA structures were disrupted or stabilized (see **Insertions are driven by the 1043-1048 cRNA structure** in main text), they remained stable in non-RdRp samples of all HA structure mutants (IN: 3.3-3.8‰ to 2.6-5.4‰, NL: 3.8-4.6‰ to 2.9-5.8‰) (**Supplementary Fig. 2**, **Supplementary Data 2**). The structure-dependent changes in insertion frequencies in RdRp-containing samples are therefore specific to RdRp-replication rather than a more general phenomenon of replication enzymes.

**Supplementary Note 5: Inconsistent single-nucleotide and homopolymer insertions in disruption mutants**

While predicted RdRp-trapping RNA structure disruption led to consistent decreases in heteropolymer and complex insertion frequencies in IN$_{RKKR}$ and NL$_{RKKR}$, its effect on single-nucleotide and homopolymer insertions was less clear (**Supplementary Data 2**). Single-nucleotide insertions did not decrease substantially with the targeted but did decrease with the full-disruption approach for both IN$_{RKKR}$ (2.4-5.1‰ to 1.1-3.8‰ or 0.1-0.3‰, respectively) and NL$_{RKKR}$ (4.6-5.3‰ to 2.7-3.6‰ or 1.7-2.0‰). Homopolymer insertions decreased with both approaches for NL$_{RKKR}$ (2.2-3.3‰ to 0.7-1.2‰ or 0.4-0.5‰), but only upon full disruption for IN$_{RKKR}$ (1.0-2.5‰ to 1.0-1.4‰ or 0.5-0.4‰).

To further investigate the inconsistent effect of structure disruption on single-nucleotide and homopolymer insertions, we designed two additional full disruption mutants for IN$_{RKKR}$, and two additional targeted disruption mutants for NL$_{RKKR}$. Different substitutions, but with similar predicted disrupted RdRp-trapping structures, had comparable and consistent effects on heteropolymer and complex insertion frequencies (**Extended Data Fig. 2c,d**, **Supplementary Data 2**), but still showed inconsistent patterns for single-nucleotide and homopolymer insertions. In IN$_{RKKR}$, single-nucleotide and homopolymer frequencies remained within a twofold range for one of the additional mutants tested, while homopolymer insertion frequencies decreased about twofold for the other (**Extended Data Fig. 2c**). In the additional NL$_{RKKR}$ targeted disruption mutants, single-nucleotide insertion frequencies either did not change or decreased about fivefold, while homopolymer insertion frequencies again consistently decreased, by four- to fiftyfold (**Extended Data Fig2d**).

**Supplementary Note 6: Differences in insertion frequencies depending on amplified RNA sense**

In all conditions using an initial cRNA template, the frequency of heteropolymer insertions was about twofold higher than observed when vRNA was used as template (8.5-10.1‰ to 15.9-20.5‰). Since heteropolymer insertions occur mainly during replication of cRNA templates (see **Heteropolymer insertions occur on cRNA** in main text), providing the RdRp with cRNA templates in abundance might lead to an increase in observed heteropolymer frequencies. On the other hand, single-nucleotide and homopolymer insertion frequencies were strongly increased when sequencing cRNA products (**Fig. 3c,d**) instead of vRNA products (**Fig. 3a,b**) (4.4-7.4‰ to 20.2-35.1‰ and 0.5-1.0‰ to 3.2-7.8‰ respectively). This seems to be linked to sequencing cRNA cleavage sites as a similar increase was observed when comparing the insertion frequencies detected in PolI-produced cRNA templates (**Fig. 3a,b**) to those in PolI-produced vRNA templates (**Fig 3c,d**) (2.5-4.1‰ to 8.2-10.6‰ and 0.0-0.1‰ to 0.3-1.7‰ respectively).

**Supplementary Note 7: H5 HPAIV precursors contain the 1043-1048 cRNA structure**

To assess the relevance of the 1043-1048 cRNA structure for LPAIV-to-HPAIV conversion in nature, we investigated whether it was present in LPAIV precursors of known H5 HPAIV outbreaks. We focused only on the eleven H5 LPAIV-to-HPAIV conversions resulting from MBCS acquisition via insertion and for which the nucleotide sequence was available[2]. Viruses from these outbreaks can be sorted into three lineages, the geographical AEO and American lineages, and a sublineage of the American lineage first isolated in Mexico in 1994 (Mx/94). Viruses from the Mx/94 lineage have evolved separately and stand out from other American H5 LPAIVs by their continuous circulation in poultry. LPAIV precursors have been identified only in three out of the eleven HPAIV outbreaks: Canada 1966, Mexico 1994, and South Africa 2006[2]. For all other outbreaks, the genetically closest available LPAIV HA identified via BLAST was used as proxy. For each LPAIV-to-HPAIV conversion event, one HPAIV representative was chosen. We then determined the transient RdRp-trapping structure predicted to form when the cRNA nucleotide 1045 was in the RdRp active site, where the lowest free energy 1043-1048 cRNA structure was identified in IN and NL (**Fig. 1b,c**). All but one of the eight putative HPAIV precursors from the AEO lineage presented the NL 1045 cRNA structure (**Extended Data Table 1**). Only the putative precursor of HPAIVs from the South Africa 1961 outbreak had a slightly less stable 1045 cRNA structure than $NL_{RETR}$. However, this LPAIV HA was relatively distant from the HPAIV HA, due to the scarcity of sequences from old H5 viruses, and all available HPAIV HA sequences of viruses from this outbreak did present the NL 1045 cRNA structure. Putative precursors of the three non-AEO HPAIV all harbored an identical predicted structure at position 1045, equivalent to the NL 1045 cRNA structure but lacking the central G-C base pair, resulting in a large decrease in stability (**Extended Data Table 1**).

**Supplementary Note 8: Deletions are promoted by A/U stretches**

Deletions were sorted into two major categories: unambiguous deletions, which can be aligned to the reference only at one position, and ambiguous deletions, which can be aligned multiple ways (usually deletions in a repetitive sequence stretch). Ambiguous deletions were aligned to the most stable predicted RdRp-trapping structure possible. They were further subdivided according to their size: single-nucleotide deletions (SND) and larger deletions. Deletion frequencies split by type for each replicate and each template are provided in **Supplementary Data 2**. All raw non-corrected deletion frequencies for RdRp-containing and background samples are provided as an interactive figure in **Supplementary Fig. 4**, alongside additional information such as alignments, predicted structures, and coverage at each position.

In all LPAIV HAs (IN$_{RETR}$, NL$_{RETR}$, SW$_{IETR}$), low frequencies of deletions were detected (<0.4‰ in RdRp-containing, <0.9‰ in background samples), but with large deletions of up to 23 nucleotides observed even at these low frequencies.

Replacing the RETR or IETR cleavage site with RKKR led to a stark increase in deletion frequencies, in both RdRp-containing and background samples, to a total of 12.9-15.4‰ for IN$_{RKKR}$ (background: 12.1-13.5‰), 11.9-12.2‰ for NL$_{RKKR}$ (background: 8.4-8.9‰) and 17.8-19.1‰ for SW$_{RKKR}$ (background: 7.0-9.3‰). Across all 21 replicates of IN$_{RKKR}$, we observed that deletion frequencies were consistent in RdRp-containing, but much more variable in background samples (**Extended Data Fig. 3c, top and bottom right panels**). In all cases, ambiguously aligned deletions represented at least 95% of all deletions observed. The SND/larger deletion frequency ratios were strongly increased in RdRp-containing samples when compared to background samples: 4.5-5.7 vs 0.9-0.9, 6.6-8.0 vs 1.0-1.5, and 5.3-8.1 vs 0.8-0.9 for IN$_{RKKR}$, NL$_{RKKR}$, and SW$_{RKKR}$ respectively, suggesting that the RdRp causes a distinct deletion profile from background deletions in RKKR-bearing templates, with more frequent SNDs. The size distribution of deletions across all IN$_{RKKR}$ replicates confirmed that SND were consistently much more common in RdRp-containing than in background samples (**Extended Data Fig. 3d, top and bottom right panels**).

### Supplementary Note 9: Deletion frequencies are not affected by predicted RNA structures

Stabilizing or disrupting predicted RdRp-trapping structures did not appreciably affect deletion frequencies, as almost all changes in frequency were below twofold. As an example, the two substitutions predicted to disrupt the 1043-1048 cRNA structure, leading to an eightfold decrease in heteropolymer insertions in $NL_{RKKR}$ (**Fig. 2c**) resulted in only slight differences in overall deletion frequencies in RdRp-containing $NL_{RKKR}$ samples (11.9-12.2‰ to 14.5-14.9‰) and in background $NL_{RKKR}$ samples (8.4-8.9‰ to 6.6-7.7‰) (**Supplementary Fig. 4**). In addition, stabilizing the 1043-1048 cRNA structure by extending it on the internal edge, which resulted in an eightfold increase in heteropolymer insertion frequencies in $NL_{RKKR}$ (**Fig. 2d**), had no effect on deletions frequencies in the RdRp-containing $NL_{RKKR}$ samples (11.9-12.2‰ to 10.4-10.7‰) or in background samples (8.4-8.9‰ to 7.9-14.8‰).

**Supplementary Note 10: Deletions by the RdRp occur mostly on cRNA**

In the unidirectional RNP replication system, deletion frequencies did not vary compared to a bidirectional system when only cRNA to vRNA replication occurred, neither for RdRp-containing (17.4-21.6‰ to 22.3-23.7‰) nor for non-RdRp samples (1.2-1.3‰ to 1.4-2.6‰). However, when cRNA to vRNA replication was blocked, deletion frequencies decreased about threefold in the RdRp-containing (18.7-20.0‰ to 6.6-7.8‰) but not in the non-RdRp samples (10.4-13.6‰ to 12.9-14.4‰) (**Supplementary Fig. 4**). Here, we observed an effect of the RNA orientation targeted for sequencing on the deletion frequencies observed in the non-RdRp samples.
