## Supplementary Table 1 for "Transient RNA structures underlie highly pathogenic avian influenza virus genesis"

### Supplementary Table 1: Primer sequences used in this study:

| Primer use | Sequence |
|---|---|
| **All constructs** | |
| PCR amplification 1 | TGTAAAACGACGGCCAGT |
| PCR amplification 2 | TAATACGACTCACTATAGGG |
| **$IN_{RETR}$** | |
| Reverse transcription | TGTAAAACGACGGCCAGTNNNNNNNNNNCAAACAGATTAGTCCTTGCAACAGG |
| 2nd strand synthesis | TAATACGACTCACTATAGGGACCTGCTATAGCTCCAAATAGTCC |
| **$IN_{REKR}$** | |
| Reverse transcription | same as $IN_{RETR}$ |
| 2nd strand synthesis | same as $IN_{RETR}$ |
| **$IN_{RKKR}$** | |
| Reverse transcription (vRNA) | same as $IN_{RETR}$ |
| 2nd strand synthesis (vRNA) | same as $IN_{RETR}$ |
| Reverse transcription B (vRNA) | TGTAAAACGACGGCCAGTNNNNNNNNNNGATTAGTCCTTGCAACAGGGCTC |
| 2nd strand synthesis B (vRNA) | TAATACGACTCACTATAGGGCCTCTATAAAACCTGCTATAGCTCCAAATAG |
| Reverse transcription (cRNA) | TGTAAAACGACGGCCAGTNNNNNNNNNNACCTGCTATAGCTCCAAATAGTCC |
| 2nd strand synthesis (cRNA) | TAATACGACTCACTATAGGGCAAACAGATTAGTCCTTGCAACAGG |
| **$IN_{RKKR}$ 1043-1048 cRNA structure disruption** | |
| Reverse transcription | same as $IN_{RETR}$ |
| 2nd strand synthesis | same as $IN_{RETR}$ |
| **$IN_{RKKR}$ with $NL_{RETR}$ 1043-1048 cRNA structure** | |
| Reverse transcription | same as $IN_{RETR}$ |
| 2nd strand synthesis | TAATACGACTCACTATAGGGACCTGCTATAGCCCCAAATAGTCC |
| **$IN_{RKKR}$ full disruption (13 substitutions)** | |
| Reverse transcription | TGTAAAACGACGGCCAGTNNNNNNNNNNGAAATCAAACAGATTAGTCCTTGCAATAAG |
| 2nd strand synthesis | TAATACGACTCACTATAGGGACTTGCTATTGTTGTAAATATTCC |
| **$IN_{RKKR}$ full disruption (12 substitutions)** | |
| Reverse transcription | TGTAAAACGACGGCCAGTNNNNNNNNNNGTGAAATCAAACAGATTAGTCATTGCAACAG |
| 2nd strand synthesis | TAATACGACTCACTATAGGGCTCTATAAAACCTGCTATAACTCCTATTAATCC |
| **$IN_{RKKR}$ disruption (7 substitutions)** | |
| Reverse transcription | same as $IN_{RETR}$ |
| 2nd strand synthesis | TAATACGACTCACTATAGGGCTCTATAAAACCTGCTATAGCTCCTATTAATCC |
| **IN-specific structure stabilization** | |
| Reverse transcription | same as $IN_{RETR}$ |
| 2nd strand synthesis | TAATACGACTCACTATAGGGTGCTATAGCTCCAGACAGTCC |
| **$NL_{RETR}$** | |
| Reverse transcription | TGTAAAACGACGGCCAGTNNNNNNNNNNCAGATAGATTAGTCCTTGCGACTGG |
| 2nd strand synthesis | TAATACGACTCACTATAGGGCTGCTATAGCCCCAAATAGTCC |
| **$NL_{REKR}$** | |
| Reverse transcription | same as $NL_{RETR}$ |
| 2nd strand synthesis | same as $NL_{RETR}$ |
| **$NL_{RKKR}$** | |
| Reverse transcription | same as $NL_{RETR}$ |
| 2nd strand synthesis | same as $NL_{RETR}$ |
| **$NL_{RKKR}$ 1043-1048 cRNA structure disruption** | |
| Reverse transcription | same as $NL_{RETR}$ |
| 2nd strand synthesis | TAATACGACTCACTATAGGGCTGCTATAGCTGCAAATAGTCC |
| **$NL_{RKKR}$ 1043-1048 cRNA structure internal stabilization** | |
| Reverse transcription | same as $NL_{RETR}$ |
| 2nd strand synthesis | TAATACGACTCACTATAGGGCTGCTATAGCCCCTCATAGTCC |
| **$NL_{RKKR}$ full disruption (10 substitutions)** | |
| Reverse transcription | same as $NL_{RETR}$ |
| 2nd strand synthesis | TAATACGACTCACTATAGGGCTACTTTAACGCTAATTAGTCC |
| **$NL_{RKKR}$ 1043-1048 cRNA structure disruption (5 substitutions A)** | |
| Reverse transcription | same as $NL_{RETR}$ |
| 2nd strand synthesis | TAATACGACTCACTATAGGGCTGCTATAGCTGTAAATAGTCC |
| **$NL_{RKKR}$ 1043-1048 cRNA structure disruption (5 substitutions B)** | |
| Reverse transcription | same as $NL_{RETR}$ |
| 2nd strand synthesis | TAATACGACTCACTATAGGGCTGCTATACAGGGAAATAGTCC |
| **$NL_{RKKR}$ 1043-1048 cRNA structure external stabilization** | |
| Reverse transcription | same as $IN_{RETR}$ |
| 2nd strand synthesis | same as $IN_{RETR}$ |
| **$SW_{IETR}$** | |
| Reverse transcription | TGTAAAACGACGGCCAGTNNNNNNNNNNTGAGGCTTGCAACTGGACTAAGA |
| 2nd strand synthesis | TAATACGACTCACTATAGGGCGATGGCTCCGAAAAGTCC |
| **$SW_{RKKR}$** | |
| Reverse transcription | same as $SW_{IETR}$ |
| 2nd strand synthesis | same as $SW_{IETR}$ |
| **$SW_{RKKR}$ 1043-1048 cRNA structure disruption** | |
| Reverse transcription | same as $SW_{IETR}$ |
| 2nd strand synthesis | TAATACGACTCACTATAGGGCGATGGCTTTGAAAAGTCC |
| **$SW_{RKKR}$ with NL 1043-1048 cRNA structure** | |
| Reverse transcription | same as $SW_{IETR}$ |
| 2nd strand synthesis | TAATACGACTCACTATAGGGCGATGGCCCCGAAAAGTCC |
